# The molecular landscape of neural differentiation in the developing *Drosophila* brain revealed by targeted scRNA-seq and a multi-informatic analysis paradigm

**DOI:** 10.1101/2020.07.02.184549

**Authors:** Nigel S. Michki, Ye Li, Kayvon Sanjasaz, Yimeng Zhao, Fred Y. Shen, Logan A. Walker, Cheng-Yu Lee, Dawen Cai

## Abstract

The *Drosophila* type-II neuroblast (NB) lineages present an attractive model to investigate the neural differentiation process. With only 16 stem cells, the type-II NB lineages generate many intermediate neural progenitors (INPs) to rapidly expand the neuron and glia pool, similar to those in the human outer subventricular zone (OSVZ). We performed targeted single-cell mRNA sequencing (scRNA-seq) in 3rd instar larval brains and created MiCV, an scRNA-seq data visualization web tool to integrate results from multiple bioinformatics analyses, display co-expression patterns of multiple genes simultaneously, and retrieve gene function and ortholog annotations. We identified novel markers that label distinct neural subsets using MiCV and subsequently *in situ* profiled them to recover the spatial information lacking in the scRNA-seq data. These new markers further enabled us to build novel neural developmental trajectories that lead to unique neuronal cell fates. Combining prior knowledge, *in silico* analyses, and *in situ* evidence, this multi-informatic investigation describes the molecular landscape of neural differentiation from a single developmental snapshot in *Drosophila*, and provides an experimental and analytical roadmap for navigating the differentiation process of more complex brains.

## INTRODUCTION

The brain is generated by a set of complex fate-specification mechanisms that birth a diverse pool of neural and glial subtypes. These mechanisms rely upon some of the approximately 1500 transcription factors found in the vertebrate genome (Zhou et al., 2017). Understanding which of these transcription factors play a role in neural fate specification remains an open area of basic research across model organisms (Bayraktar and Doe, 2013; Homem and Knoblich, 2012; Soldatov et al., 2019; Zhong et al., 2018). In particular, untangling the interplay of intrinsic (cell-specific) and extrinsic (global, spatial) fate patterning mechanisms remains particularly challenging, especially in the complex and large vertebrate brain.

*Drosophila melanogaster* represents a model organism that recapitulates features of vertebrate neurogenesis. Unlike the abundant type-I neuroblasts (NB, neural stem cells), the 16 type II NBs in the *Drosophila* brain adopt a neurogenesis process that is directly analogous to that observed in mammalian cortical development (Homem and Knoblich, 2012). During development, each type II NB undergoes repeated asymmetric cell divisions to generate an NB and a sibling progeny that acquires a progenitor identity (i.e. intermediate neural progenitor, INP). Each INP undergoes limited rounds of asymmetric cell division to re-generate and to produce a ganglion mother cell (GMC), which divides once more to become two neuron(s) and/or glial cell(s). Along this NB-INP-GMC-neuron maturation process, cells express a well-defined cascade of transcription factors that mark these cell differentiation stages (Ren et al., 2017; Syed et al., 2017). In parallel, INPs born in each division cycle may express a unique cascade of transcription factors that contribute to the generation of different neural progenies (Bayraktar and Doe, 2013). It is highly plausible that the combination of these two transcription factor cascades alongside a third molecular axis, which defines unique NBs (i.e., each NB generates a distinct lineage), brings about the generation of a highly diverse neuronal pool (**Fig. 1A**).

**Figure 1.**
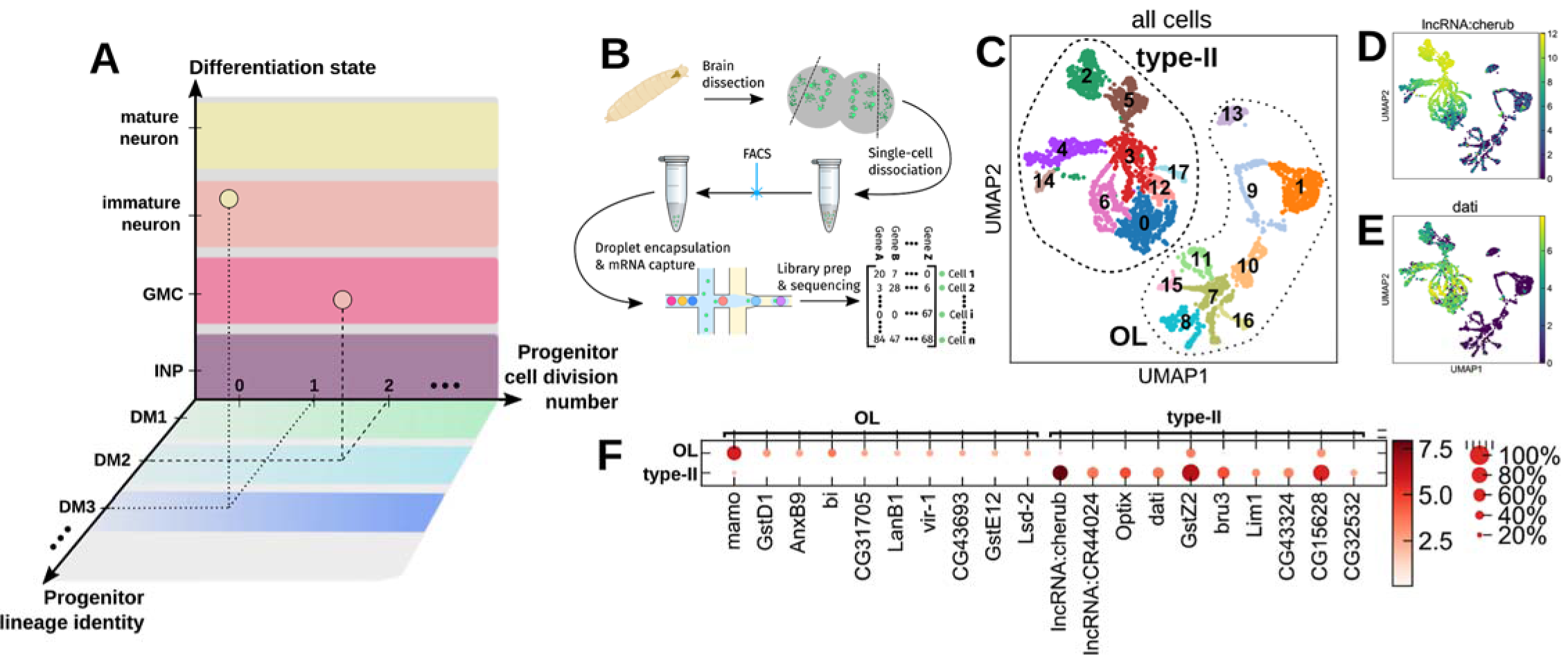
Drosophila type-II neuronal fate specification model, experiment overview, and *in silico* dissection of the optic lobe and type-II derived cells. **(A)** A diagram of the major axes that determine cell “state” in this work. Each sequenced cell is defined in part by factors that are specific to the lineage identity, intermediate progenitor cell division number, and differentiation state. Untangling these axes enables us to more fully reconstruct the neural fate specification landscape of type-II neurogenesis at a specific developmental time point. **(B)** Late stage 3rd instar larval brains were dissected and optic lobes were partially removed before enzymatic and mechanical single-cell dissociation. Fluorescent reporter-positive cells were FACS selected. A single cell mRNA library prepared by a droplet-based scRNA-seq platform (10X Chromium) is subsequently sequenced and aligned to the *Drosophila* transcriptome. **(C)** After initial QC filtering, cells were plotted in the first 2 dimensions of a UMAP projection. Color represents an automatic cluster assignment by the Leiden algorithm (resolution = 0.5). **(D-E)** Expression of the long non-coding RNA *cherub* and the transcription factor *dati* are known to be exclusive of the optic lobe in 3rd instar larvae. Groups of cells that lack expression of these genes are likely optic lobe cells that also express *Gal4* under the control of the R9D11 fragment of the *erm* promoter. **(F)** Separating the putative type-II/optic lobe cells into two groups and performing logistic regression analysis reveals genes that are up-regulated between the two.

The advent of high throughput single-cell mRNA sequencing (scRNA-seq) technologies has enabled researchers to broadly investigate the mRNA expression landscape of hundreds of thousands of cells (Macosko et al., 2015; Ziegenhain et al., 2017). Coupled with a wide variety of analytical tools (Butler et al., 2018; Wolf et al., 2018), researchers can make hypotheses about the number of unique cellular subtypes in the brain (Cocanougher et al., 2019; Saunders et al., 2018), what the functions of these subtypes might be (Ren et al., 2019), and what subtypes might arise together along a common developmental pathway (Cao et al., 2019; Qiu et al., 2017; Soldatov et al., 2019). While such “cell atlas” style scRNA-seq datasets effectively characterize the transcriptomes of a majority of cells from a region of interest, cell populations that are classically clustered together (through *in situ* and/or functional analyses, for example) may not be identified by blind *in silico* cluster analysis (Kiselev et al., 2019). In addition, broad scRNA-seq studies often do not take advantage of the extensive collection of genetic labelling tools that can highlight classically clustered cell populations, enabling them to be studied in greater detail. For instance, a targeted approach to scRNA-seq is required if we are to confidently and efficiently describe nuanced developmental systems, such as the specification of unique neural subtypes derived from the type-II NB lineages of *Drosophila*, where inclusion of non-type-II derived cells (making up the majority of the fly brain) would introduce overwhelming noise and confound our analysis.

In the type-II NB lineages of *Drosophila*, we set out to broadly classify the molecular factors that define the neural progenies of dividing INPs along three key fate-patterning axes, i.e., differentiation state, division number, and progenitor lineage (**Fig. 1A**) using targeted scRNA-seq. We created a long-living fluorescent reporter to brightly label the type-II progenies at the 3rd instar larval stage and FACS sorted them in preparation for 10X Chromium scRNA-seq (**Fig. 1B**). We subsequently recovered transcriptomes containing 11,187 genes from 3942 cells. Through an iterative process of cell clustering, marker gene analysis, pseudotime analysis, and *in situ* validation, we identified genes that vary in expression along all three neural fate-patterning axes mentioned above. These genes include markers that globally define the INP, GMC, and neuron differentiation stages in most NB lineages. Further *in silico* analysis suggested molecular factors that are uniquely expressed in subpopulations of INPs, GMCs, immature and mature neurons. Subsequent *in situ* mRNA staining recovered the spatial relationship of these molecular factors, which clarified the cell division number and NB lineage specificity. We finally identified novel markers that exclusively label distinct neural subsets. These new markers further enabled building novel neural developmental trajectories that lead to unique neuronal cell fates. Our multi-informatic approach to targeted scRNA-seq experimental design and analysis provides a roadmap for navigating the differentiation process of complex brains.

## RESULTS

### Type-II neuroblast derived cells are uniquely identified from the mixed optic lobe cell population using descriptive quality control metrics and clustering

To perform *targeted* scRNA-seq, we brightly labeled the type-II NB progenies with a long-lasting fluorescent reporter. We created an UAS-hH2B::2xmNG reporter fly, in which two copies of the mNeonGreen (2xmNG) fluorescent protein are fused to the C’-terminus of the human histone 2B protein (hH2B). This leverages the expression of multiple copies of a bright fluorescent protein alongside the slower turn-over rate of the histone protein (Tumbar et al., 2004). To validate labeling fidelity, we compared the expression pattern of UAS-hH2B::2xmNG to UAS-IVS-myr::tdTomato under the control of the R9D11-Gal4 driver, in which Gal4 is active in type-II INPs and a small subset of medial optic lobe (OL) cells in larval brains (Weng et al., 2010). While the overall labeling patterns of these two UAS transgenes were similar, hH2B::2xmNG labeled more cells to form larger clusters than the membrane-targeted IVS-myr::tdTomato clusters, which indicated that carry-over hH2B::2xmNG labeled more post-mitotic neurons (**Suppl. Fig. 1**). Finally, the bright nuclear mNG labeling enabled reliable FACS selection for targeted 10x Chromium scRNA-seq (**Fig. 1B**, and detailed in **Methods**).

Subsequently, we projected the dimension-reduced scRNA-seq data onto a 2D UMAP plot and overlaid the counts of all genes, unique transcripts (UMI), and mitochondrial genes as part of routine scRNA-seq quality control (**Suppl. Fig. 2**). When overlaying the hH2B::2xmNG reporter transcript counts to the 2D UMAP, we found that mNG transcripts were expressed non-uniformly, with pockets of cells expressing the hH2B::2xmNG transcript at a level that was nearly 4 orders of magnitude greater than others in the dataset (**Suppl. Fig. 3A**). To examine whether this non-uniform expression pattern reflects true biological variance, we performed *in situ* RNA staining for *mNG* using the HCRv3 protocol (Choi et al., 2018) and imaged the native mNG fluorescence to compare the relationship of mNG transcripts and proteins (**Methods**). We found that each of the type-II clusters indeed expresses a high level of *mNG* transcripts in only a small subpopulation of cells near the tip of each lineage (**Suppl. Fig. 3B-D**). This spatial localization, coupled with co-expression of *mNG* transcripts with *D* in *CycE+* cells (data not shown) leads us to conclude that the R9D11 enhancer fragment’s expression is tightly restricted to newly-born INPs and their daughter GMCs, emphasizing the need for long-living reporters for investigation of neural subtypes derived from the type-II NBs.

**Figure 2.**
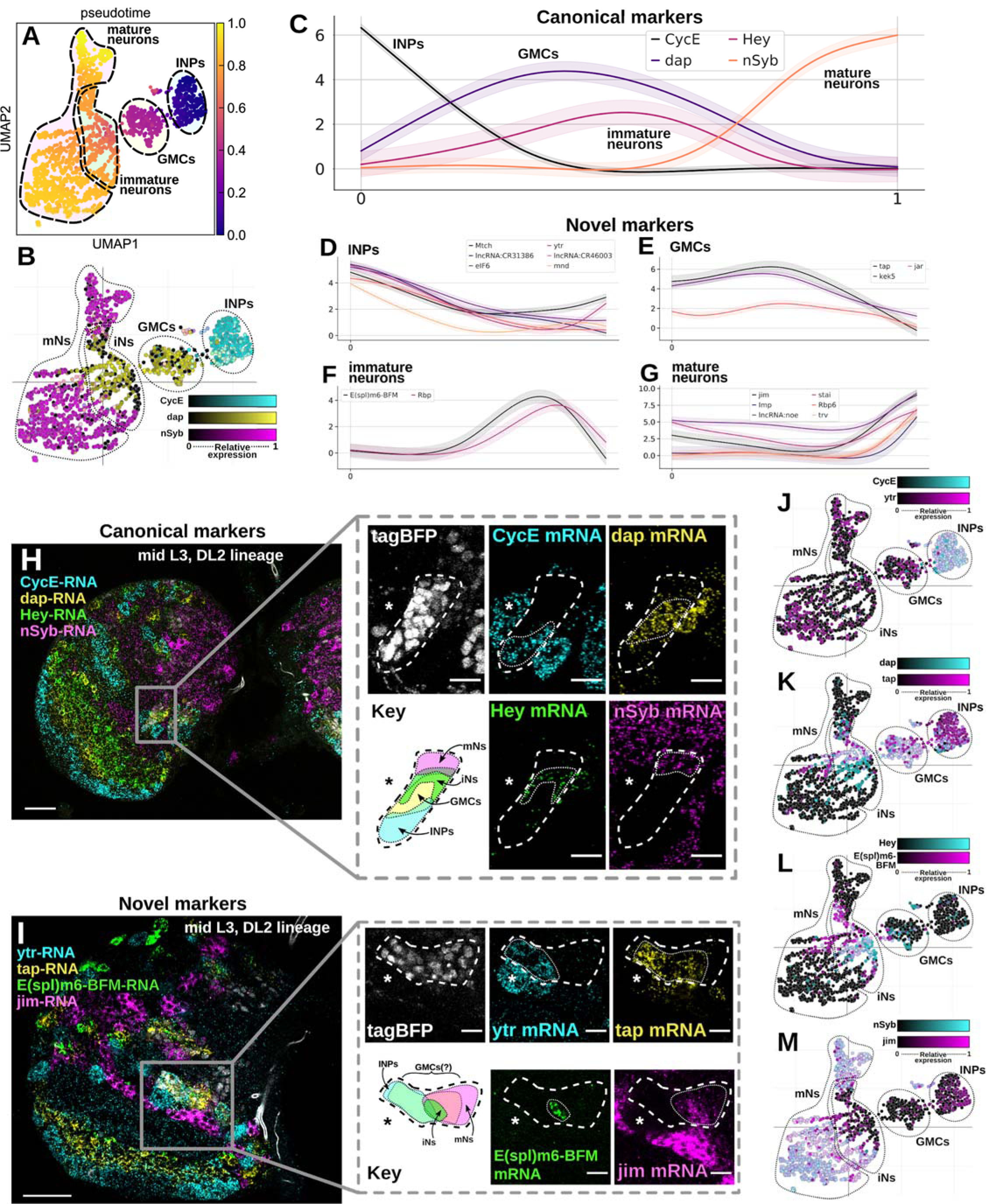
Pseudotime analysis reveals signature genes that vary along the cell differentiation axis. **(A)** Pseudotime analysis (performed using scanpy and the palantir Markov-chain-based pseudotime algorithm) establishes a global ordering of cells along the differentiation state axis. **(B)** A multi-color UMAP expression plot generated by the MiCV web tool shows the expression patterns of 3 canonical marker genes for the classical INP, GMC, and mature neuron states. **(C)** The pseudo-temporal expression pattern of 4 genes that are known markers for the 4 major differentiation states. **(D-G)** Pseudo-temporal expression patterns of groups of marker genes that do not have known functions associated with cellular differentiation state. These genes vary strongly along the pseudo-temporal axis in patterns similar to the known marker genes plotted in (C). **(H, I)** HCRv3 *in situ* mRNA staining images for both known (H) and novel (I) differentiation state marker genes in single z-slices of the DL2 lineage of mid 3rd instar larval brains. UAS-hH2B::2xtagBFP is driven under the control of R9D11-Gal4 and marks the type-II lineages. Magnified inserts show individual spectral channels and diagrams of the gene expression pattern. Asterisks denote the location of the putative type-II NB. Thick dashed lines denote the boundaries of the tagBFP labeled type-II NB progenies. Thin dotted lines denote the boundaries of type-II progeny cells expressed indicated mRNAs. **(J-M)** Multi-color UMAP expression plots illustrate the expression pattern of the canonical and novel marker genes from (H) and (I), respectively. Scale bars: 30 μm in overviews of (H, I), 10μm in insets of (H, I).

**Figure 3.**
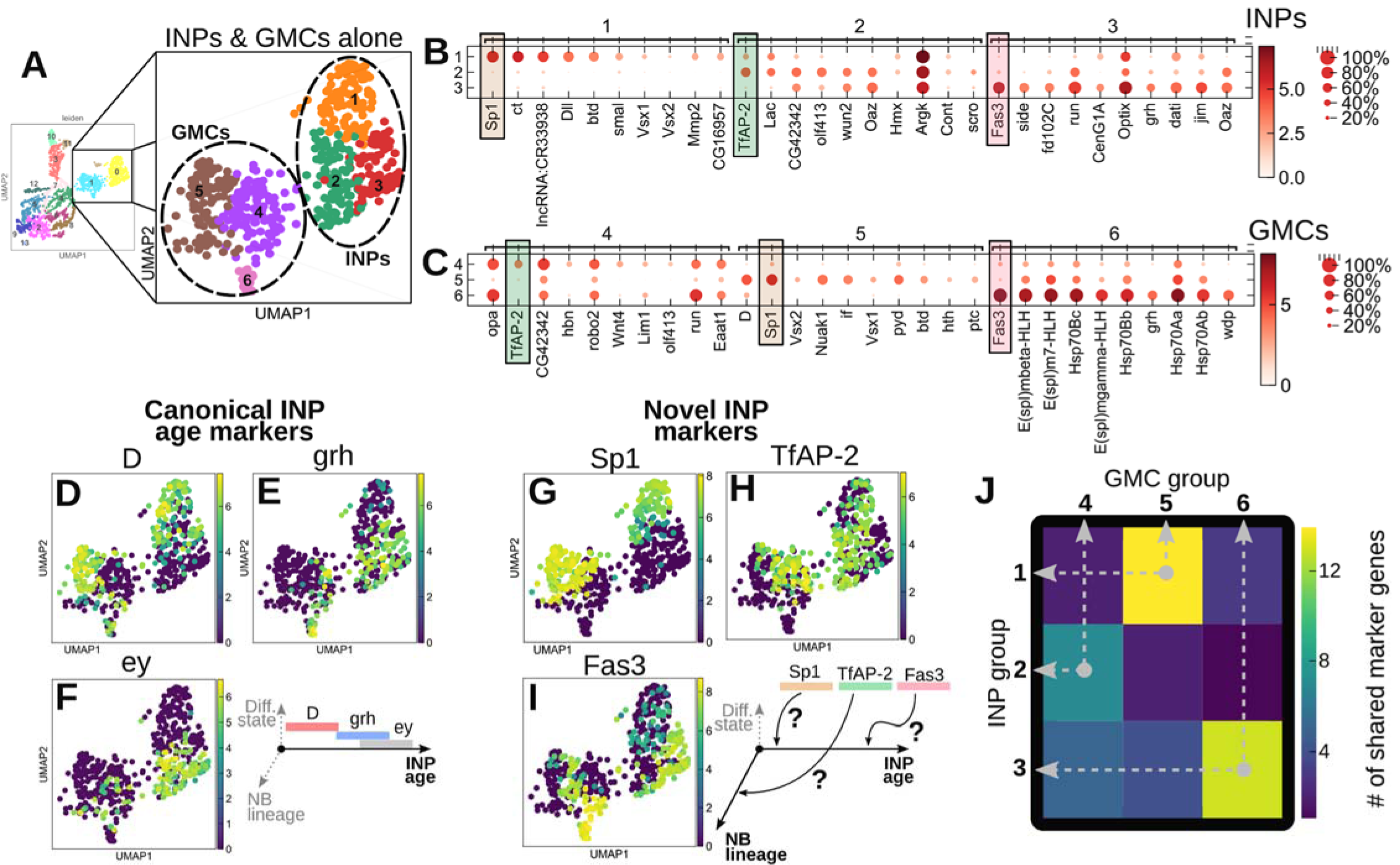
Sub-clustering of INPs and GMCs reveals transcription factors beyond the canonical *D*-*grh*-*ey* transition that vary along a combination of the NB lineage and INP division number patterning axes. **(A)** Right panel, low resolution clustering reveals INPs and GMCs to be in cluster 0 and 1, respectively. Left panel, higher resolution clustering on separated INPs and GMCs further divides them into 3 subclusters each. **(B, C)** Marker gene analysis revealed that mostly transcription factors specific INP and GMC subclusters, respectively. The color-coded genes highlight a strong correspondence between corresponding INP and GMC subclusters. **(D-F)** Expression UMAP plots of the well-established temporally-varying INP genes *D, grh*, and *ey*, respectively. *D* appears to separate cleanly at the mRNA level in the INPs of our dataset, however, *grh* and *ey* are broadly co-expressed. **(G-I)** Expression UMAP plots of the INP/GMC cluster-specific genes *Sp1, TfAP-2*, and *Fas3*, which are found to correlate INP subclusters 1, 2, and 3 to GMC subclusters 5, 4, and 6, respectively. **(J)** A correlation plot shows the number of top 100 marker genes that are shared between each INP and GMC subcluster. This simple similarity metric indicates a hypothesis that cells in GMC subclusters 5, 4, and 6 are the direct progenies of cells in INP subcluster 1, 2, and 3, respectively.

To further ensure the specificity of our analysis to type-II cells, we performed an *in silico* filtering to exclude the optic-lobe cells that are also labeled by R9D11-Gal4 (Boone and Doe, 2008). Using the expression of two optic lobe exclusive genes as a guide (*lncRNA:cherub* and *dati*; see *in situ* expression patterns from (Landskron et al., 2018; Schinaman et al., 2014), respectively), we identified two groups of cells which we labelled as the putative type-II (*cherub*+/*dati*+) and OL (*cherub*-/*dati*-) cells, respectively. The split of these two genes cleanly segregates our scRNA-seq data into two large subsets that are spatially separated in the initial UMAP projection (**Fig. 1C-E**). To identify other potential marker genes of OL vs. type-II cells, we performed a logistic regression-based marker gene analysis (Ntranos et al., 2018) comparing these two major groups against one another (**Fig. 1F**). The transcription factor *mamo* is the most upregulated in the putative OL cells when compared to the putative type-II cells. This upregulation is interesting as *mamo* is most well known for its role in meiosis, being required for the development of fully functional eggs (Mukai et al., 2007). Additionally, we found that *Optix* is upregulated in a subset of the putative type-II cells but not in the medial OL cells. This is consistent with *in situ* staining of *Optix* transcripts (Gold and Brand, 2014), which show that *Optix* is specifically expressed in the lateral part of the third instar larval OL. The subset of type-II cells expressing *Optix* are likely those from lineages DM1-3, based on *in situ* expression data (Gold and Brand, 2014). Our scRNA-seq data also indicates that *Optix* co-expresses in *D* or *bsh* expressing type-II cells (data not shown), which have been previously identified as part of the DM1-3 lineages (Bayraktar and Doe, 2013; Boone and Doe, 2008). From this *in silico* filtering process that relies largely on *in situ* expression data from the literature, we confidently separated the type-II derived cells from optic lobe cells that were also captured in our scRNA-seq experiment. Only these type-II derived cells were carried forward for our downstream analysis.

### Pseudotime analysis describes the continuous differentiation stages of type-II derived cells

Knowing that the *R9D11-*hH2B::2xmNG reporter specifically labels type-II progenies from INPs to maturing neurons, we aimed to first align each cell along the continuous cellular differentiation state axis (**Fig. 1A**). We expected this would reveal the most prominent underlying structure of our data because, in the case of type-II neurogenesis, all cells will similarly transition through the INP, to GMC, to immature, to mature neuron differentiation states. Using the Markov chain-based pseudotime analysis algorithm *Palantir* was a natural choice as Markov chains describe discrete transitions that occur randomly based upon a continuous probability distribution (Setty et al., 2019). Given a properly chosen starting cell, *Palantir* aligns cells in our scRNA-seq data based upon the path of fewest transcriptomic changes propagating from the starting cell.

Cells expressing high levels of the INP markers *CycE* and *D* are good candidate starting cells for *Palantir* (Bayraktar and Doe, 2013; Yang et al., 2017). To easily identify these cells from the UMAP plot, we built a <u>M</u>ulti-informatic <u>C</u>ellular <u>V</u>isualization web tool (MiCV) to allow displaying the single cell co-expression pattern of multiple genes in the 2D/3D UMAP plots. Furthermore, users can conveniently select a subset of cells for specific analysis, such as picking the starting cell(s) for *Palantir*, by combining mouse-click selections from the parallel plots generated by MiCV (**Methods**). We overlaid the pseudotime result onto the reprojected 2D UMAP plot that only included type-II NB derived cells. Based on the expression of known marker genes (**Suppl. Fig. 4**), we predicted INP, GMC, immature, and mature neuron clusters (**Fig. 2A**, dash lines). Interestingly, these cell maturation state clusters aligned well with the pseudotime arrangement. For example, using MiCV, we displayed the single cell co-expression pattern of *CycE, dap*, and *nSyb* (**Fig. 2B**), which are known to distinguish the INP, GMC/immature neuron, and mature neuron states, respectively, and found their UMAP positions matched well with their pseudotime alignments (**Fig. 2A**).

**Figure 4.**
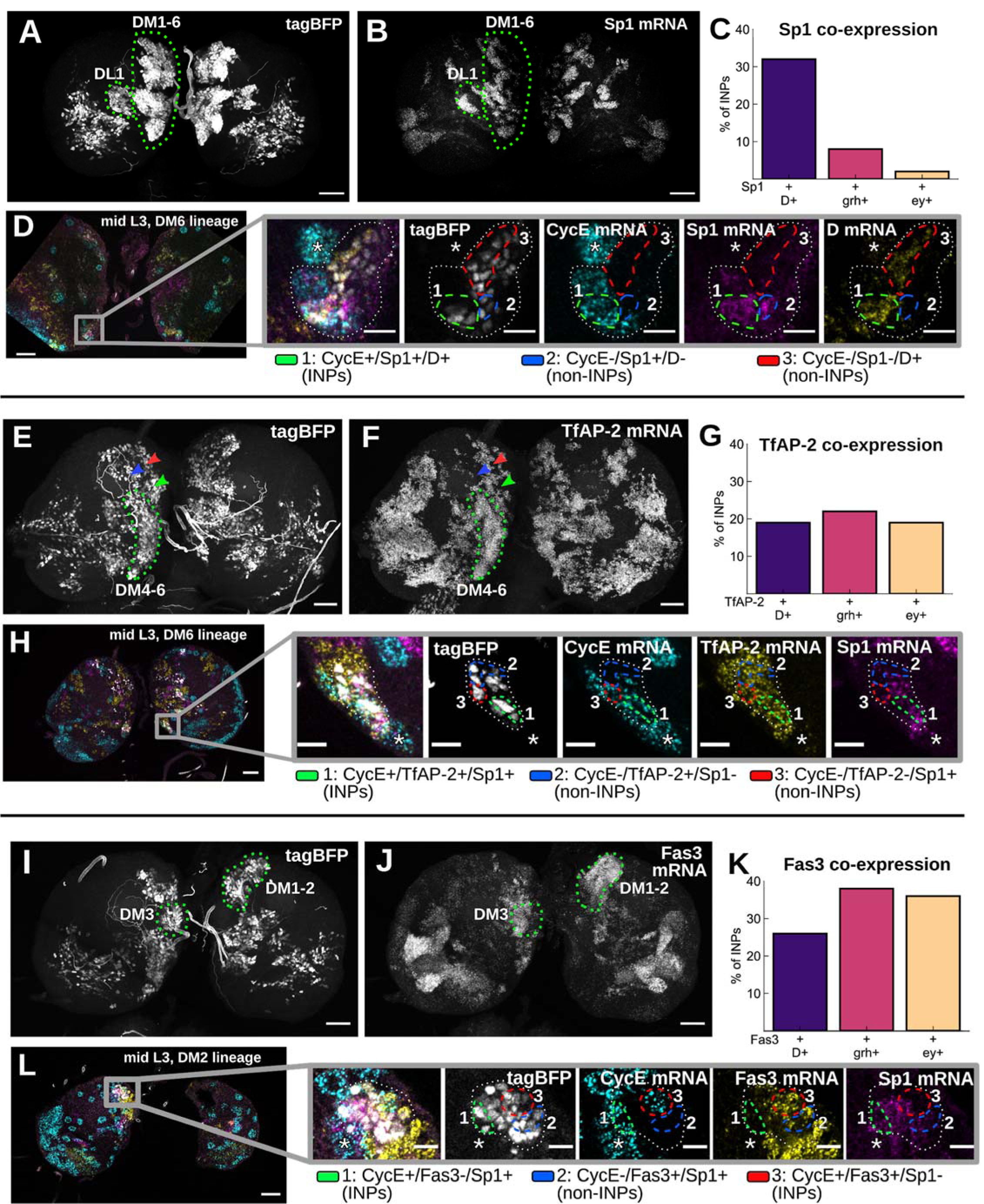
Sp1, TfAP-2, and Fas3 are each expressed by INPs of specific NB lineages. **(A, B)** Maximum Z-projections (45μm thick) show tagBFP fluorescence and *Sp1* mRNA HCR staining in an L3 larval;;R9D11-Gal4/UAS-H2B::tagBFP fly brain, respectively. Green dotted lines indicate the expression of *Sp1* mRNA in all type-II NB derived lineages except for DL2. **(C)** *Co-*expression quantification of *Sp1* with *D, grh*, and *ey* in all INPs (n=284). **(D)** HCR staining showcases the expression patterns of *Sp1* and *D* mRNAs in lineage DM6. Dashed lines highlight region 1 of INPs that co-express *Sp1* and *D* mRNA, region 2 of non-INP cells where *Sp1* mRNA alone is detected, and region 3 of non-INP cells where D mRNA alone is detected. White dotted lines denote the DM6 lineage boundary. Asterisks denote the position of the DM6 neuroblast. **(E, F)** Maximum Z-projections (45μm thick) as in (B,C), with *TfAP-2* mRNA HCR staining. Within the type-II NB lineages, *TfAP-2* mRNA is highly expressed in cells belonging to DM 4-6 (dotted lines) and possibly DL1. Though some expression is seen nearby to DM1-3, *TfAP-2* is not expressed in tagBFP+ cells belonging to those lineages (arrowheads). **(G)** Co-expression quantification of *TfAP-2* with *D, grh*, and *ey* in all INPs (n=284). **(H)** HCR staining showcases the expression patterns of *CycE, Sp1* and *TfAP-2* mRNAs in lineage DM6, where we can find *TfAP-2* expressed in *CycE*+ INPs that have *Sp1* expression (green dashed lines) or not (red dashed lines), as well as in *CycE*-progeny cells (blue dashed lines). **(I, J)** Maximum Z-projections (45μm thick) as in (B,C), with *Fas3* mRNA HCR staining. Within the type-II NB lineages, *Fas3* mRNA is highly expressed in cells belonging to DM 1-3 (dotted lines). **(K)** Co-expression quantification of *Fas3* with *D, grh*, and *ey* in all INPs (n=284). **(L)** HCR staining showcases the expression patterns of *CycE, Fas3*, and *Sp1* mRNAs in lineage DM2. We find a clear expression of *Fas3* in both INPs and their progeny in NB lineage DM2, where we can find *Fas3* expressed in *CycE*+ INPs that have *Sp1* expression (green dashed lines) or not (red dashed lines), as well as in *CycE*-progeny cells (blue dashed lines). Scale bars: 30 μm in (A, B, E, F, I, J) and in overviews of (D, H, L); 10 μm in insets of (D, H, L).

To describe the dynamics of gene expression across pseudotime, and thus the differentiation process, we fit a gene expression trend line to each gene detected in our scRNA-seq dataset using *Palantir*. Indeed, we found that the expression peaks of four marker genes, i.e. *CycE* for INPs (Yang et al., 2017), *dap* for GMCs (Lane et al., 1996; de Nooij et al., 1996), *Hey* for a subset of the transient immature neuronal state (Monastirioti et al., 2010), and *nSyb* for maturing neurons (Deitcher et al., 1998), aligned in this exact differentiation order along the calculated pseudotimeline (**Fig. 2C**). Hence, we can use the relative expression levels of these genes to approximate the boundaries of the continuously changing differentiation states (**Fig. 2A**, dashed lines) in pseudotime. Subsequently, we performed gene expression trend clustering (from Palantir) to screen novel putative marker genes whose expression trend matched one of the four known marker genes’ (**Fig. 2D-G**). Independently, we used a marker gene-based differentiation state scoring (Wolf et al., 2018) strategy to separate these differentiation stages and found similar sets of marker genes (**Suppl. Fig. 4**). Interestingly, many of the putative marker genes do not have any known function related to neural differentiation. Further pathway analysis and gene manipulation studies will be needed to explore their exact roles in type-II neurogenesis.

Nonetheless, we profiled the *in situ* expression patterns of some putative marker genes we identified in this analysis. We consider this to be essential not only for the validation of the *in silico* discovery but also for the selection of candidates to explore their novel functions. We first synthesized HCRv3 probes against the canonical makers *CycE, dap, Hey*, and *nSyb* transcripts (**Methods**) and used these probes to investigate their expression pattern in the type-II NB derived cells using our novel reporter fly. As predicted, these genes form largely non-overlapping expression patterns in the larval brain (**Fig. 2H**, left panel). When zooming into the DL2 type-II lineage (**Fig. 2H**, right panel), we found that *CycE* transcripts were expressed in large neuroblasts as indicated by the large cell bodies (**Fig. 2H**, right panels, asterisk) and in smaller tagBFP positive cells as a marker for replicating INPs. As predicted, *dap, Hey*, and *nSyb* transcripts expressed in bands of cells that were sequentially positioned away from the neuroblast (**Fig. 2H**, right panels, dashed lines). Next, from the gene expression trend clustering result (**Fig. 2D-G**), we selected a set of four candidate novel markers and performed similar HCR *in situ* mRNA profiling. The *in situ* results suggest that *ytr, tap, E(spl)m6-BFM*, and *jim* transcripts express in unique patterns (**Fig. 2I**, right panels) and the co-expression MiCV plots indicate that these markers largely overlap the canonical makers in the respective cells (**Fig. 2J-M**). In particular, *E(spl)m6-BFM*, and *jim* were expressed almost exclusively in immature neurons and maturing neurons, respectively (**Fig. 2L-M**). However, while the putative INP marker *ytr* expressed in 96% of all the INPs, it also expressed in 37% of GMCs and 38% of maturing neurons (**Fig. 2J**). This observation indicates that *ytr* broadly expresses in INPs and that its expression may be selectively maintained in a subset of GMCs and their progeny neurons similar to a canonical transcription factor *D* (detailed below). The putative GMC marker *tap* appears to express in subsets of INPs and approximately half of the immature neurons (**Fig. 2K**). This suggests that *tap* may be a gene that defines one daughter immature neuron apart from the other during their mother GMC’s terminal cell division (see further discussion below).

Though many genes that trend along the differentiation state axis are potentially interesting, we highlight here the gene *E(spl)m6-BFM*, a member of the *Notch*-responsive subgroup of the “enhancer of split” family of transcription factors (Lai et al., 2000). This family of proteins is responsible for regulating a variety of developmental processes (Maier et al., 1993) and their group’s function in balancing the self-renewal of differentiation in the type-II neuroblasts of *Drosophila* has recently been described (Li et al., 2017). However, the specific function or restricted spatial expression of *E(spl)m6-BFM* in the developing larval brain has not been established. Based on our analysis, *E(spl)m6-BFM* may mark *all* the cells in a transient immature neuronal state, which comes about directly after the mother GMC’s terminal cell division. This is in contrast to *Hey*, the currently known immature neuron marker, which is upregulated in only one of the two daughter neurons of this terminal GMC division (Monastirioti et al., 2010) and activates in a *Notch*-dependent manner. Indeed, *E(spl)m6-BFM* is expressed in both *Hey*+ cells and *Hey*-cells that have similar pseudotime values (data not shown). Interestingly, the novel marker gene *tap* is expressed exclusively in the half of immature neurons that are *Hey*- (magenta cells in **Fig. 2K** *vs* cyan cells in **Fig. 2L**). Similar to *E(spl)m6-BFM, Rbp*, a protein known to be functionally required for synaptic homeostasis and neurotransmitter release (Liu et al., 2011; Müller et al., 2015), is also upregulated only in this immature neuronal subset (data not shown). Further study will be desired to understand why either of these genes undergo a burst of expression in the *immature* neuronal state, and to establish their functional roles at the protein level.

### INP and GMC sub-clustering enables the identification of novel maturation pathways that are convolved with the canonical *Dichaete, grainy-head, eyeless* transition

Having used pseudotime analysis to define the major differentiation states in the type-II neurogenesis process, we next characterized the cellular heterogeneity within these states using automated scRNA-seq clustering analysis. Such analysis may or may not obviously reflect previously established models of cell type differentiation/diversity, especially when this diversity could refer to any of/all the axes of cell type differentiation (**Fig. 1A**). Nonetheless, we performed Leiden clustering (Traag et al., 2019) with a low resolution (0.6) and overlaid the result on the reprojected UMAP (**Fig. 3A**, left). We found that cluster 0 and 1 included 284 and 259 cells, which correspond to the INP and GMC populations in the above-mentioned pseudotime analysis, respectively. Subsequently, we took these putative INP and GMC cells and found they could be subdivided into three groups of INPs and three groups of GMCs in using a clustering resolution of 0.6 (**Fig. 3A**, right).

To discover which genes distinguished each subcluster, we performed logistic regression-based marker gene analysis and plotted the top 10 genes that defined the INP (**Fig. 3B**) or GMC subclusters (**Fig. 3C**). We found that this clustering result does not reflect the canonical *Dichaete, grainy-head, eyeless* transition, which has been indicated to sequentially express in young to old INPs over the course of division cycles (Bayraktar and Doe, 2013). *D* expression was rather specific in 74% of subcluster 1 INP cells and in 78% of subcluster 5 GMC cells, while only expressing in fewer than 28% of other subcluster cells (**Fig. 3D**). On the contrary, *grh* and *ey* expressions are intermingled in the other subclusters (**Fig. 3E** and **3F**, respectively). Interestingly, we found that *Sp1, TfAP-2*, and *Fas3*, the top marker genes in this clustering analysis, not only expressed in segregated subclusters, but also marked both INP and GMC subclusters (**Fig. 3G, 3H**, and **3I**, respectively). We suspected that the GMC subclusters specified by these genes might be the direct progenies of the INP subclusters that carry over the *Sp1, TfAP-2*, and *Fas3* transcripts. We subsequently counted the number of top 100 marker genes that were shared between each of the INP and GMC subclusters. The correlation plot strongly suggests that GMC subcluster 5, 4, or 6 is likely the progeny of INP subcluster 1, 2, or 3, respectively (**Fig. 3J**).

The choice of clustering resolution can be somewhat arbitrary, and the 3 subclusters for INPs and for GMCs here may represent a surface level of INP patterning that can be further broken down into more subtypes. Although we attempted clustering INPs and GMCs at a significantly higher resolution and performed similar marker gene analysis, the low cell counts in each cluster made assessment of more subclusters less statistically meaningful (data not shown). Therefore, we opted to focus on *in situ* validation of the marker genes identified at the 3-subcluster resolution in follow up experiments.

### The transcription factor *Sp1* is expressed in young INPs throughout the DM1-6 and DL1 lineages and marks a unique neural progeny

We first aimed to *in situ* profile the transcript expression of the marker gene *Sp1*, a Cys2His2-type zinc finger transcription factor that has recently been shown to be necessary, alongside its partner gene *btd*, for the specification of type-II neuroblasts (Álvarez and Díaz-Benjumea, 2018). We reasoned that this, along with the apparent coexpression of *Sp1* with *D* in the INPs of our scRNA-seq dataset (**Fig. 3D, 3G**, respectively), would imply that *Sp1* may be broadly expressed in young, newly-matured INPs of most type-II NB lineages. We synthesized HCRv3 probes against *Sp1* and *D* transcripts (**Methods**) and validated their specificity using gene-trap reporter flies (**Suppl. Fig. 5**). When accessing their expression patterns in the type-II NB derived cells using our novel reporter fly, we found that *Sp1* mRNA was expressed prominently in all type-II lineages with the exception of DL2 (**Fig. 4A, 4B**). On the contrary, *D* mRNA expressed prominently in DM1-3, and in much smaller subsets of cells in lineages DM4-6 (data not shown), which is consistent with previous observations (Bayraktar and Doe, 2013).

**Figure 5.**
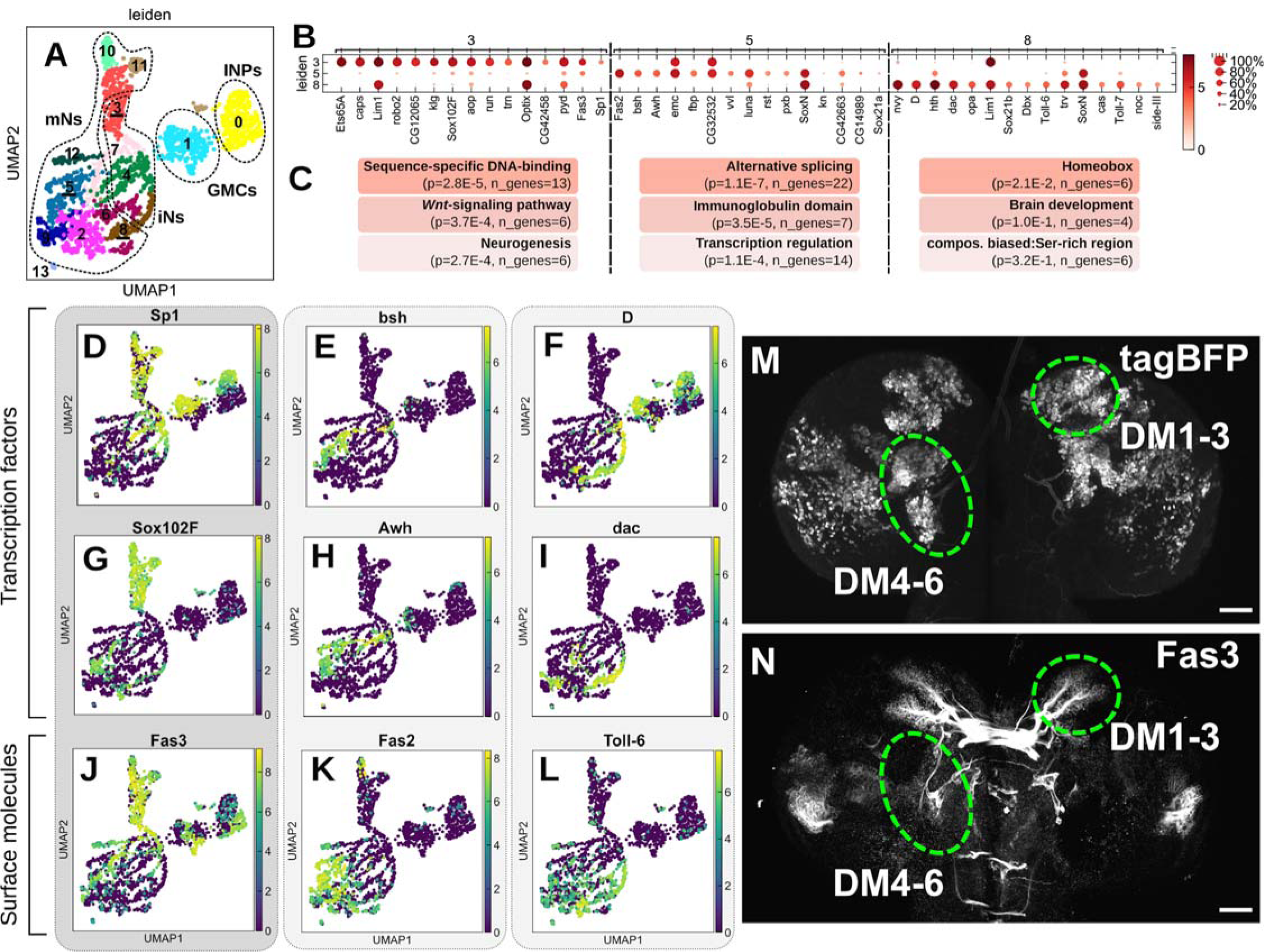
Transcription factors and surface molecules predominantly mark the late 3rd instar neuron subtypes. **(A)** Automatic Leiden clustering (resolution = 0.6) of the type-II scRNA-seq data, with putative neural subtypes 3, 5, and 8 outlined, representing the *Sp1, bsh*, and *D*+ neural progenies, respectively. **(B)** Marker gene detection for the three selected neural subtypes showing the top 15 marker genes as identified using the t-test_overestim_var function in scanpy. **(C)** The top gene ontology (GO) functional annotations for the top 100 marker genes for cells in each of clusters 3, 5, and 8, respectively (p-values are Benjamini corrected; n_genes refers to the number of marker genes annotated with the respective GO term). **(D, E, F)** Log-fold expression values of *Sp1, bsh*, and *D*, respectively, showing three unique neural lineages are marked by these three transcription factors. **(G, J)** Log-fold expression values of the transcription factor *Sox102F* and the cell surface molecule *Fas3* that mark the *Sp1*+ neural progeny. **(H, K)** Log-fold expression of the transcription factor *Awh* and surface molecule *Fas2* that mark the *bsh*+ neural progeny. **(I, L)** Log-fold expression of the transcription factor *dac* and surface molecule *Toll-6* that mark the *D*+ neural progeny. **(M, N)** Maximum Z-projections show tagBFP fluorescence and *Fas3* antibody staining in an L3 larval;;R9D11-Gal4/UAS-H2B::tagBFP reporter fly brain, respectively. It appears that neurons from DM1-3 that produce commissure-crossing axons are prominently labeled by *Fas3*, whereas neurons from DM4-6 are largely unstained. Scale bars: 30 μm in (M, N).

Our scRNA-seq data indicates that while *Sp1* co-expressed with *D* in more than 30% of INPs (**Fig. 4C**), 8% and 16% of all INPs are *Sp1*+/*D*- and *Sp1*-/*D*+, respectively. To validate the presence of these INP populations *in situ*, we used our HCR protocol to co-stain *Sp1* and *D* mRNA using our novel reporter fly (**Fig. 4D**). Here, we show that, for instance in the DM6 lineage, an *Sp1*+, *D*+ INP progeny can be identified directly adjacent to cells where either *Sp1* or *D* is exclusively expressed (**Fig. 4D**, enlarged box). Furthermore, we overlaid *Sp1* or *D* expressions on the UMAP plot, and found that these two transcripts continue to express in maturing neurons of two exclusive subsets (detailed below). This is consistent with a previous study, which found that the *D* expressing young INPs specifically give rises to *D* expressing neurons (Bayraktar and Doe, 2013). Therefore, we hypothesize that *Sp1*+/*D*+ INPs may transition to *Sp1* or *D* exclusively expressing INPs, which give rise to distinct neuronal subtypes. To specify whether *Sp1* protein is expressed in neurons, we labeled the type-II progenies with a membrane-bound tdTomato (R9D11-CD4::tdTomato) to visualize neuron’s characteristic axonal projections and coupled with an Sp1::GFP reporter line. We show that, for instance, the DM3 lineage generates many neurons that form a *tdTomato*+ neurite bundle are also *GFP*+, which indicates the generation of *Sp1*+ neural progeny (**Suppl. Fig. 5B**).

Next, we wondered whether *Sp1* is like *D* that expresses strictly in young INPs. We quantified our scRNA-seq data and found that *Sp1* coexpressed with the two canonical late INP markers *grh* and *ey* only in a small subset of INPs (**Fig. 4C**). Taken together, these data support the hypothesis that *Sp1*, much like *D*, is expressed broadly in INPs with low division numbers and that these INPs are responsible for producing a neural progeny similarly marked by *Sp1* expression that is distinct from the *D*+ neural progeny.

### The transcription factor *TfAP-2* and cell adhesion molecule *Fas3* are each expressed in INPs of specific type-II neuroblast lineages

We next characterized the spatial expression patterns of *TfAP-2* and *Fas3*, selected markers for the other two major putative INP subtypes identified in our low-resolution clustering (**Fig. 3**). We generated HCR probes against mRNA of *TfAP-2* and *Fas3* in a similar manner to *Sp1* and probed their expression in reporter flies in order to identify which type-II NBs generate their respective INP subsets. Unlike *Sp1*, however, *TfAP-2* and *Fas3* transcripts are expressed much more broadly across the brain and are not restricted to the type-II lineages (**Fig. 4F, 4J**).

Within the type-II progenies, *TfAP-2* mRNA appeared to be expressed prominently in INPs of the DM4-6 lineages as well as a subset of their downstream progeny (**Fig. 4E, 4F**, green outline). However, we did not observe strong *TfAP-2* expression in any other lineage, implying that expression of this marker is primarily lineage restricted (**Fig. 4E, 4F**, arrowheads). Interestingly, *TfAP-2* co-expressed in fewer *D*+ but many more *grh*/*ey*+ INPs than *Sp1* does in our scRNAseq data, which indicates that *TfAP-2*+ INPs have likely undergone some cell divisions before expressing this marker gene (**Fig. 4C** vs **4G**). Although *TfAP-2* expresses in fewer lineages than *Sp1*, our scRNA-seq data (not shown) and *in situ* profiling (**Fig. 4H**) showed that these two genes do indeed co-express in cells belonging to those few lineages. *TfAP-2* plays broad roles in development (Monge et al., 2001), but in the context of the central brain it has been shown to play a role in developing and maintaining the neural circuitry required for night-sleep in adult flies (Kucherenko et al., 2016). Consistently, we found in our scRNAseq data that *TfAP-2* expressed in a subset of neurons that are distinct from the *Sp1+* or *D*+ population (data not shown). *TfAP-2*’s expression in neurons is distinct from the previously identified late INP progeny genes *grh* and *ey*; the latter two were not found in neurons in our scRNA-seq data (data not shown). *TfAP-2* (*ap-2*) is significantly orthologous to the human transcription factors TFAP2A/B (Flybase curators, 2019), and its role in sleep can be traced back to *C. elegans* (Turek et al., 2013). Taken together, this would imply that at least this particular role for *TfAP-2* in the central brain may be evolutionarily conserved and that the neurons generated by *TfAP-2*+ INPs in the DM4-6 lineages may play a role in night-sleep circuit maintenance.

Based on our *in situ* RNA staining, *Fas3* mRNA was found to express most prominently in the INPs of DM1-3 (**Fig. 4I, 4J**). Similar to *TfAP-2*, our scRNAseq data suggests that *Fas3* co-expressed in fewer *D*+ but many more *grh*/*ey*+ INPs than *Sp1* does, which indicates that *Fas3* INPs have likely undergone some cell divisions before expressing this marker gene (**Fig. 4C** vs **4K**). Again, our scRNA-seq data (not shown) and *in situ* profiling (**Fig. 4L**) showed that *Fas3* and *Sp1* co-express in a significant fraction of cells. *Fas3* is interesting as a marker gene for INPs as it is *not* a transcription factor but rather a membrane-bound, homophilic cell adhesion molecule that plays a strong role in synaptic targeting and axonal guidance in a subset of neurons in the central and peripheral nervous systems (Kose et al., 1997; Snow et al., 1989), along with cell adhesion-mediated morphological development throughout the entirety of the fly (Wells et al., 2013). Why *Fas3* would be expressed so strongly in a subset of INPs is unknown.

### Transcription factors and surface molecules, not neural functional genes, are the primary neural subtype markers of the type-II neural lineages at the 3rd instar larval stage

With the low resolution (0.6) clustering, our scRNA-seq data already showed a much greater subtype diversity in neurons (12 clusters) than in GMCs or INPs (1 cluster each) (**Fig. 5A**). We performed logistic-regression based marker gene analysis on these specific clusters and identified the top 100 marker genes for each cluster that are most uniquely expressed, the top 10 of which are plotted in **Suppl. Fig. 6**. Such automatic cluster analysis enables *de novo* identification of the *Sp1*+, *D*+, and *bsh*+ neural lineages that compose significant proportions of cell clusters 3, 8, and 5, respectively (**Fig. 5B**). Subsequently, we analyzed the top 100 marker genes using the DAVID Functional Annotation Tool (Huang et al., 2009a, 2009b) in order to identify sets of genes that form functionally associated groups based on associated gene ontology (GO) terms. We identified the first GO term from the top three highly enriched functional groups and find that these terms indicate that transcription factors and surface molecules are predominant markers for these three (**Fig. 5C**), as well as all other neural subsets (data not shown). We focused on these three populations because in the (Bayraktar and Doe, 2013) study, *bsh* was found to express in a non-overlapping subset of neurons that do not express *D* in the young INP progeny. The same study also specified that there are more young INP derived neurons are *Bsh*- and *D*-. Therefore, we hypothesize that *Sp1* contributes to the specification of the remaining non-identified young INP derived neuron population. Indeed, our scRNAseq data shows that *Sp1, D*, and *bsh* were expressed in three distinct maturing neuron populations (**Fig. 5D, 5F**, and **5E**, respectively).

**Figure 6.**
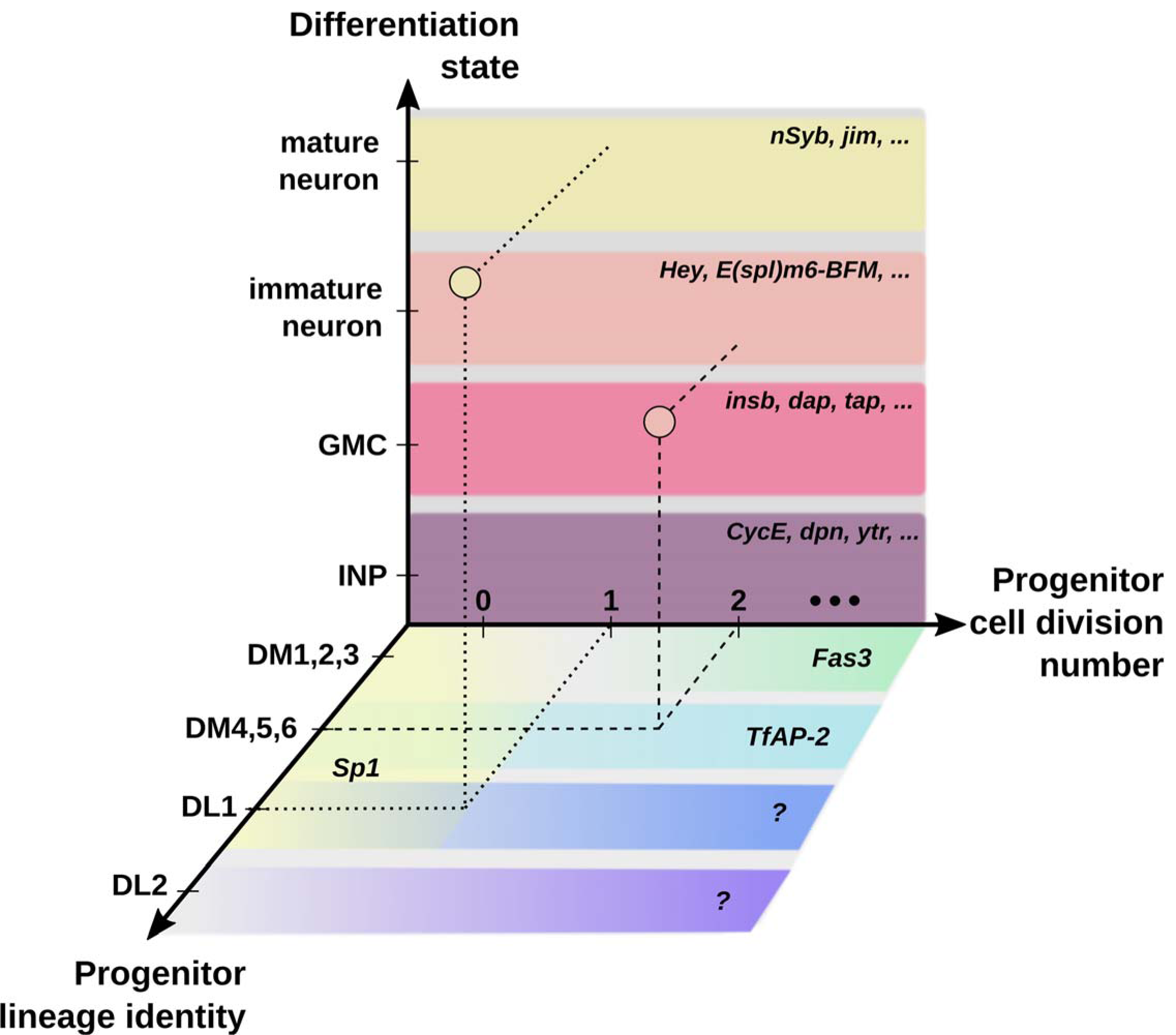
A Drosophila type-II neuronal fate specification model illustrates the complex molecular network that determines the neural differentiation process. Despite its small scale and apparent simplicity, the complex interplay of molecular factors that vary along the differentiation state, lineage identity, and progenitor cell division number axes are responsible for determining the fate of each cell derived from the type-II neuroblasts of *Drosophila*. In this diagram, some of the most prominent molecular factors from the literature or identified and validated in this work are shown to occupy different domains along these three axes. Continuing analysis and *in situ* validation will enable us to continue to fill in the blanks and develop a more complete roadmap of the type-II neurogenesis process at a single developmental timepoint. However, data from multiple developmental timepoints, alongside methods for overlaying lineage identity onto scRNA-seq data, will be required to accomplish this in a more holistic and high-throughput manner.

This *in silico* analysis permits rapid identification of novel transcription factors that potentially belong to the same regulatory pathways that specify neuronal fate. For example, selected from the specific marker gene list, the transcription factors *Sox102F, dac*, and *Awh* are highly co-expressed with neurons expression *Sp1, D*, and *bsh*, respectively (**Fig. 5G, 5I, 5H**, respectively). Distinct surface molecules are also differentially expressed in different subsets of neurons, which may indicate their roles in forming functionally distinct circuits (**Fig. 5J-5L**). Among them, *Fas3* appears to co-express in a large proportion of *Sp1* neurons, regardless of their low degree of co-expression in the INP and GMC stages (compare **Fig. 5D** vs **5J**). To validate that Fas3 protein is indeed expressed in neurons as a functional molecule, we used a Fas3 antibody to stain our novel type-II lineage reporter fly and found that it labels neurons in the DM1-3 lineages that form neurite bundles across the commissure (**Fig. 5M, 5N**). It is plausible that the expression of *Fas3* in INP may play a role in enabling some of the neural progenies of DM1-3 to either form these axonal bundles or for them to find their final targets across the commissure early on in the neural maturation process. We conclude from this analysis that groups of transcription factors and surface molecules, as opposed to groups of genes with other terminal neuronal functions, are what most prominently specify these neural lineages at the late 3rd instar stage.

## DISCUSSION

### Drosophila type-II neural lineages as a model system to study complex neurogenesis processes

To enable the brain’s complex functions, vastly diverse neuronal types need to be rapidly generated at a very large scale during development. As an example, in a short period, the mammalian cortex expands vertically and horizontally to become a multi-cell layer structure that covers a large surface area (Greig et al., 2013). To reveal how neural stem cells populate the developing brain, efforts have been made to identify cell types and their lineage relationships. For instance, focuses on neurodevelopment in mouse (Habib et al., 2017; Han et al., 2018; Ponti et al., 2013; Saunders et al., 2018; Soldatov et al., 2019), human brain tissues (Habib et al., 2017) and the developing human prefrontal cortex (Zhong et al., 2018) revealed intermediate stem cells (and critical genes involved) as an important mechanism for rapid cortical expansion. Underlying this rapid and diverse differentiation process is the constant change of gene expression profiles in all cells. However, the molecular mechanisms that lead to functionally distinct neurons in the mammalian brain remain challenging to describe in detail. This is because, on the one hand, neuronal fate determination involves many genes, and on the other hand, neural progeny cells originating from distinct lineages undergo rapid migration, which leads their intermingling nature in space.

Although they are the minority (8 stem cells per hemisphere) in the *Drosophila* central brain, the *Drosophila* type-II neural lineage has a neurogenesis process analogous to the mammal’s rapid cortical expansion (Homem and Knoblich, 2012). The type-II neural lineage structure includes a neuroblast stem cell (NB) that divides many times to self-renew and generate an intermediate progenitor cell (INP) in each division. Each INP continues to divide many times to self-renew and in each division, generate a ganglion mother cell (GMC), which divides one more time to generate two terminal cells of neuronal or glial fate. Compared to their mammalian counterparts, the *Drosophila* type-II neural lineage has the advantage of being non-migrating in the larval stage. With proper labeling, type-II progeny cells of the same lineage can be identified as a segregated cell cluster. Importantly, the cells’ spatial relationship within a cluster serves as a considering factor when determining the age and maturation stage of these cells (Boone and Doe, 2008; Homem and Knoblich, 2012). The small stem cell pool and mammal-like lineage composition make the *Drosophila* type-II neural lineage an attractive model to study the complex brain development process. In addition, many important genes and signaling pathways are conserved throughout evolution (Homem and Knoblich, 2012; Mariano et al., 2020; Ogawa and Vallender, 2014), which makes revealing the molecular mechanisms of *Drosophila* type-II neuron differentiation a meaningful primer to study the human analogs in the outer subventricular zone.

### Summary of this work

In this work, we take advantage of targeted single cell transcriptome analysis to advance our understanding of the *Drosophila* type-II neuron differentiation process. After initially separating the transcriptomes of the type-II neuroblast derived cells from those labeled in the optic lobes, we show that pseudotime analysis techniques can be used to define a maturation axis and extract novel putative marker genes that specify the INP, GMC, immature neuron and mature neuron differentiation stages. Broadly expressed, not limited to the type-II NB progenies, these novel marker genes of different maturation stages indeed form intersectional patterns that represent the spatial organization of the neurogenesis progress in the larval brain. Compared to previous antibody-based and gene manipulation-based screening strategies, scRNA-seq data permits a high-throughput assessment of the whole gene expression profile to rapidly identify candidate genes for functional study. For instance, in the past, *Hey* has been shown to mark one of the two immature neurons derived from the final cell division, and its role is exclusive as an inhibitor of *Notch* signaling in this immature neuron (Monastirioti et al., 2010). From our scRNAseq analysis, *E(spl)m6-BFM*, a member of the enhancer-of-split family of transcription factors (Lai et al., 2000), and *Rbp*, a rim-binding protein responsible for synaptic homeostasis and neurotransmitter release (Liu et al., 2011; Müller et al., 2015) are almost exclusively up-regulated in *only* the transient immature neuronal differentiation state directly after GMC division. These two marker genes can be used to guide the exploration of *Hey*-immature neurons in future studies. Functional knock-outs of these two genes will be critical to understanding their function in newly-born neurons as it pertains to their maturation and any early functional role they may play in the developing brain.

Further higher-resolution clustering of the INP and GMC cells identified transcriptomically correlated subclusters between these two stages, which supports the idea that parallel maturation transitions happen at the same developmental time point. However, scRNA-seq data alone cannot distinguish whether these parallel transitions are due to the co-existence of earlier and newly born INPs in all NB lineages or due to the intrinsic differences among NB lineages. We therefore *in situ* profiled the marker genes selected from the scRNA-seq selected candidates and restored their missing spatial information that indicates the maturation stage as well as the NB lineage identity. In addition, combined with prior knowledge, whether a marker gene is expressed in younger or earlier born INPs can also be speculated. Our findings conclude that *Sp1* is expressed in the young INPs of nearly all NB lineages, whereas *TfAP-2* and *Fas*3 express in older INPs belonging to specific NB lineages. Interestingly, we found that *Sp1* and *TfAP-2* expressed not only in neural progenitors but also in maturing neurons. These transcription factors seem to intermingle with the *D*/*grh*/*ey* cascade in the INP stage but eventually differentiate into completely exclusive neuron populations. Finally, higher-resolution clustering of neurons in our scRNAseq dataset revealed that transcription factors and surface molecules are predominant markers for distinct neuronal subtypes at the 3rd instar larval stage. This implies that most neurons of the type-II NB progenies have not started to gain their differentiated functions at this stage of development.

During data analysis, we realized the importance of being able to conveniently display and integrate analytical results generated by multiple bioinformatics pipelines. Aiming to allow biologists to efficiently explore scRNA-seq data without the need of any coding, we developed MiCV, an open-source web tool that integrates multifaceted bioinformatics analysis and generates publication-ready plots requiring only a few mouse clicks and simple keyboard inputs. We found that MiCV’s novel function of plotting multiple genes in the same UMAP graph greatly helped us to efficiently investigate their expression relationship and dynamics at the single cell level. Together with an automated gene function curating feature, MiCV is highly relevant to developmental biology and cell heterogeneity studies. Combining MiCV exploring and *in situ* mRNA imaging, we discovered many transcription factors and surface molecules that potentially play important roles in generating neuronal subtypes in an NB-specific, INP-specific, or function-specific manner. These novel discoveries helped us to gain a comprehensive understanding of the molecular landscape along all three major neural developmental axes that define a cell’s progenitor lineage identity, progenitor cell division number, and differentiation state (**Fig. 6**). This model provides a general guidance for biologists to disentangle the differentiation process in complex systems beyond the *Drosophila* brain.

### Challenges and opportunities

We sequenced approximately 1800 cells that were neurons originating from 8 *Drosophila* type-II neuroblast lineages (16, if we assume no symmetry across the two central brain lobes). With low-resolution clustering, we identified 10 molecularly distinct neural subtypes. Increasing the clustering resolution just a bit higher we could identify more than 20 that are still distinct (data not shown). Similarly, as we show with the INPs/GMCs in our dataset, a low-resolution clustering can often mask the cellular diversity that is present in the system. As we know that each type-II neuroblast generates approximately 38 INPs throughout their developmental lifespan (Bayraktar et al., 2010; Bello et al., 2008), the presented clustering in this paper only captures *part* of the INP diversity. One straightforward thought is to increase the number of sequenced single cells so that higher clustering resolution may eventually reveal even the most subtle differences between each of the hundreds of INPs in the type-II system. However, as transcription factor cascades involved in INP division/maturation intertwined with those involved in NB specification or general cell differentiation stages, we expect that the INP heterogeneity can be untangled somewhat using a higher clustering resolution but still fails to provide us with a coherent view of the complex lineage, maturation, and differentiation landscape we are attempting to characterize. This challenge of deconvoluting the INP maturation, NB lineage, and differentiation state axes highlights the need for a holistic, integrated approach to experimental design and subsequent bioinformatic analysis.

The data we have presented here were collected at a single developmental time-point (late third instar), but we know that type-II neurogenesis precedes and continues after this stage. Repeating these scRNA-seq experiments at the second instar and early pupal time-points will enable us to describe more completely in what order molecularly-defined neural subsets are generated. However, this stronger introduction of the developmental time axis will not immediately make clear which combination of neuroblast lineage, INP maturation, and developmental time specifies each neural subset, as even at a single developmental time-point this specification is already difficult to untangle. Using recently developed analytical techniques to “stitch” these multi-time-point datasets together (Lin et al., 2019; Tran and Bader, 2019) will be advantageous to align all the cells along a unified developmental time axis. However, the introduction of single-neuroblast lineage barcoding techniques will be necessary to deconvolve data from this axis. Genetic constructs based around CRISPR-Cas9 (Raj et al., 2017; Spanjaard et al., 2018) and the Cre/Lox system (Kalhor et al., 2018; Pei et al., 2017; Weber et al., 2016) have been developed for this purpose, although which exact lineage was labeled by a particular barcode was still unknown. The introduction of a spectrally unique barcode for each neuroblast lineage, in a similar vein to the recently developed *Bitbow* lineage tracking strategy (Li et al., 2020; Veling et al., 2019), would be advantageous as they can provide direct *in situ* evidence for neuroblast lineage identity.

## DATA AVAILABILITY

Raw sequencing data generated in this study will be available from the Gene Expression Omnibus (www.ncbi.nlm.nih.gov/geo/).

## CODE AVAILABILITY

Jupyter notebooks used for scRNA-seq analysis are available upon request. The source code for the MiCV web tool is available at www.github.com/cailabumich/MiCV. A web server with preloaded datasets, including the one reported here is available at www.micv.works.

## Supporting information

Resource Table

## ACKNOWLEDGMENTS

We thank all members of the Cai lab who contributed to the discussion and revision of the manuscript. NSM acknowledges support from the National Institute of Health (NIH) 1T32EB005582. FYS acknowledges the support from NIH 1F31NS11184701. DC acknowledges support from the University of Michigan (CDB IDEA Awards in Stem Cell Biology, MCubed2.0) and Michigan Economic Development Corporation (Mi-TRAC). C-YL acknowledges support from the NIH 1R01NS107496.

## AUTHOR CONTRIBUTIONS

NSM and DC conceived of the project and designed experiments with critical inputs from YL and C-YL. FYS developed the HCR protocol for larval brain staining. NSM, YL, KS, and YZ performed experiments. NSM and LAW performed data analysis. NSM designed and developed the MiCV software. NSM and DC wrote the manuscript with critical insights from YL, LAW, and C-YL. DC initiated and supervised the project.

## CONFLICTS OF INTEREST

NSM is the founder of MiOmics Inc. (MI, USA). All other authors declare no conflicts of interest.

## METHODS

### Material availability

Further information and requests for resources should be directed to the lead contact: Dawen Cai.

Fly lines generated in this study include the [;;UAS-hH2B::2xmNG] and [;;UAS-hH2B::2xtagBFP] lines which will be deposited to the Indiana Bloomington Drosophila Stock Center.

Plasmids generated in this study include the pMUH-20xUAS-hH2B::2xmNG/2xtagBFP-p10pA plasmids used to generate the aforementioned fly lines and will be deposited to the Addgene plasmid repository.

HCR probes used in this study were designed and synthesized by Molecular Instruments (Los Angeles, CA, USA) and their exact sequences are the intellectual property of the aforementioned company. The target RNA sequences we provided to Molecular Instruments to design probe sets (20 probes, in 10 pairs) against are provided in the Key Resources Table, along with their lot numbers which can be used to request the exact probes used in this study.

### *Drosophila* husbandry

Flies were reared at 25°C on standard CT medium with a 12h/12h light/dark cycle.

For FACS selection of type-II derived cells for scRNA-seq, [;;R9D11-Gal4] (BD40731) virgin female flies were crossed to male [;;UAS-hH2B::2xmNG] (this study) flies in vials prepared with fresh yeast paste to promote mating. F1 progeny were collected at approximately the late 3rd instar stage, as larvae are crawling up the vial walls to prepare for pupation. No selection was made based on larval sex.

For IHC and HCR experiments, [;;R9D11-CD4::tdTom] (BD35847) virgin female flies were crossed to male flies of the following genotypes: [;;Sp1::EGFP] (BD38669), in vials prepared with fresh yeast paste to promote mating. Alternatively [;;R9D11-Gal4] (BD35847) virgin female flies were crossed to male [;;UAS-hH2B::2xtagBFP] (this study) flies in a similar manner. F1 progeny were collected at approximately the late 3rd instar stage, as larvae are crawling up the vial walls to prepare for pupation. No selection was made based on larval sex.

### Dissociation and FACS selection of type-II derived cells for scRNA-seq

[;;R9D11-Gal4/UAS-hH2B::2xmNG] larvae (n=20) were rinsed and their brains dissected using dissection scissors and forceps at the late L3 stage (wandering larvae) in ice cold Rinaldini’s solution. These brains were subsequently transferred to a poly-L-lysine coated coverslip that was immersed in Rinaldini’s solution, attaching only the VNC to coverslip and leaving the central brain lobes unattached. These brain lobes were then further dissected using a tungsten needle by inserting the needle into each brain lobe at approximately the midpoint of the lobe and moving the needle laterally. This process removed a lot but not all the cells on the lateral portions of each brain lobe, which includes the developing optic lobe. The remaining OL cells were later excluded from our final scRNA-seq dataset using known marker genes (detailed above).

Dissected brains were transferred to a DNA low-binding 1.5mL tube in 30µL of dissection liquid (Rinaldini’s solution) using a p200 pipette equipped with a siliconized p200 tip that was cut and flame-smoothed approximately 1/4 of the way up the tip. The siliconized tips are lower-binding and make it less likely for brains to stick to them. Cutting the tip and smoothing the opening makes it easier for the brains to move into the tip. The 1.5mL tube was pre-filled with 50µL of fresh, cold Rinaldini’s solution, and upon transfer of the brains, 10µL of 20mg/mL papain, 10µL of 20mg/mL type-I collagenase, and 1µL of 15µM ZnCl were added to the tube, bringing the total reaction volume to 100µL. The tube was closed and mixed gently by flicking, then incubated on a heat block at 37°C for 1hr. During this incubation, the tube was flicked for mixing at 10min intervals, flicking the tube until the brains are visibly disturbed into the solution.

After the 1hr incubation, 2µL of 100µM E-64 solution was added to the mixture to stop the papain digestion. To break down the apparent intact brains, the mixture was triturated at a ∼1 Hz frequency for 30 times using a p100 pipette set to 70µL and equipped with an uncut p200 siliconized tip. After the first 5 triturations, the brains should be seen largely dissociated to the naked eye. Further triturations break down the brain completely into single-cell suspensions including the VNC, which is quite resilient to dissociation.

After trituration, the cell suspension was diluted with 400µL Schneider’s media + 10% FBS which further quenches the enzymatic digestion and stabilizes the cells. 1µL of DRAQ5 DNA stain (Thermo Fisher Scientific Inc.) was added to label cells apart from debris generated in the dissociation process.

The sorting-ready cell suspension was transferred to a 5mL plastic FACS snap-cap tube on ice. Cells from non-Gal4 driver brains were dissociated in a similar manner and were sorted first on a Sony MA900 FACS machine to set the gate for using DRAQ5 to separate DNA containing cells from debris and set the gate for non-mNG expressing cells.

Sorted cells were captured in a DNA low-binding 1.5mL tube pre-filled with 100µL of Schneider’s media + 10% FBS. Cells were spun down at 400x g for 4 minutes and the solution volume was reduced to 40µL before resuspending by gentle pipetting with a p200 siliconized pipette tip. 5µL of this suspension was removed to count cells using an epifluorescence microscope by plating them in a single well of a 96 well plate, pre-filled with 45µL of Schneider’s media + 10% FBS. The rest of the cells were transported on ice to the University of Michigan Advanced Genomics Core and approximately 10,000 cells were loaded for 10X Chromium V3 sequencing following the manufacturer’s instruction.

### HCRv3 *in situ* mRNA staining of L3 larval brains

We adapted with only minor changes from protocols described in the original third generation HCR paper (Choi et al., 2018). In brief, late-stage third instar larvae were dissected in room-temperature (RT) PBS as previously described and transferred to a 500µL tube containing PBS on ice. Brains were washed once in PBS for 1min standing, then washed in 4% RNase-free PFA at RT with 0.5% Tween-20, nutating for 20min. Brains were then washed twice with RNase-free 0.5% PBSTween for 20min each, nutating. Brains were then washed with 200µL of HCR amplification buffer at 37°C, nutating for 1hr. HCR probes (Molecular Instruments) were added to a final concentration of 5nM, and the sample was incubated at 37°C overnight, nutating. After this incubation, brains were washed 2x in HCR washing buffer at RT for 30min each, nutating. Brains were then incubated in 200µL amplification buffer at RT for 30min, nutating. 2.5uL of each imager hairpin (with attached dyes) was independently raised to a temperature of 95°C for 90sec in a thermocycler then snap-cooled to 4C immediately. 2µL of each hairpin was then added to the brains and incubated overnight at RT, nutating. Finally, brains were washed 2x with 2X SSCT at RT for 30min each, nutating, then once again with 2X SSC at RT for at least 10 minutes, nutating. Brains were subsequently mounted on a coverslip coated with poly-L-lysine that is submerged with Prolong Diamond mounting media (Thermo Fisher Scientific Inc) for imaging.

### HCR probe design

Sequences provided to Molecular Instruments for HCR probe design were constructed by identifying the largest contiguous sequence present across all unique transcripts for each of our mRNAs of interest. As the information on the relative expression of individual isoforms of each transcript is in general not readily available, this provided for the highest possible detection probability at the expense of transcript-isoform specificity.

### IHC staining of L3 larval brains

Brains were dissected in PBS and fixed in 4%PFA + 0.5% TritonX-100 (Triton) at RT for 20min, nutating. Brains were rinsed 2x with PBST (0.5% TritonX-100) at RT, then washed 1x with 0.5% PBSTriton at RT for 30min, nutating. For primary antibody staining, brains were incubated in Starting Block + 0.5% Triton at RT for 30min. Antibodies were then added and the brains were incubated at 4C overnight, nutating. Brains were then washed as described above followed by incubating in Starting Block + 0.5% Triton at RT for 30min. Secondary antibodies were added and brains were incubated at RT for 2hr, protected from light. Brains were finally rinsed 2X in PBST (0.5% TritonX-100) at RT for 1min each, then washed 2X in PBS for 30min. Brains were subsequently mounted as described in the HCR section above.

Antibodies were diluted as the following: Mouse-anti-Fas3 1:50 (DHSB), Rat-anti-dpn (1:1000) (C-YL lab), Donkey-anti-Mouse (AF488) 1:500 (Jackson ImmunoResearch Laboratories, Inc.), Donkey-anti-Rat (AF647) 1:500 (Jackson ImmunoResearch Laboratories, Inc.).

### scRNA-sequencing and mapping

Approximately 10,000 type-II derived cells were used as input to a single channel of a 10X Chromium v3 chip. The mRNA was subsequently reverse transcribed, amplified, and prepared for sequencing on an Illumina NovaSeq-6000 chip (University of Michigan Advanced Genomics Core). The library was sequenced for a total of 385M paired-end reads with 28bp for the cell barcode and UMI and 110bp for cDNA inserts.

Reads were subsequently mapped using both Cell Ranger (for initial analysis) and STAR-solo (for our final analysis, with mNG added to the genome) (Dobin et al., 2013) to the Drosophila genome assembly provided by ENSEMBL, build BDGP6 (2014-07). Cell Ranger identified 3942 cells with a mean read count of 97,604 reads/cell, corresponding to a median of 1829 unique genes/cell and 6044 UMIs/cell.

### scRNA-seq filtering, dimensionality reduction, and clustering analysis

The downstream scRNA-seq analysis was performed using scanpy (Wolf et al., 2018), and our analysis was formalized into the MiCV web tool generated in this work (www.micv.works). In brief, cells were filtered by requiring between 200-4100 unique genes/cell (to exclude debris and some doublets) and genes were filtered by requiring at least 2 cells to express it at greater than 1 UMI/cell. UMI counts were normalized to a total sum of 1e6 UMIs/cell and subsequently log-transformed by calculating ln(1+UMIs) for each gene for each cell. The top 2000 highly variable genes were identified using the cell-ranger method (Zheng et al., 2017) and these genes were used to perform a principal component analysis (PCA, n=50pcs), neighborhood identification (k=20), and finally a UMAP projection (2D). Clusters were identified using the Leiden algorithm (Traag et al., 2019) or an optimized version of the Louvain algorithm (Blondel et al., 2008), with varying clustering resolutions. For most of the type-II only UMAP projections displayed in this work, the clustering resolution was 0.6, with 1 being a standard default (and higher numbers leading to more granular clustering of cells). Marker genes were identified using logistic regression analysis, implemented in scanpy.

### scRNA-seq pseudotime trajectory inference

Pseudotime trajectories were generated using palantir (Setty et al., 2019). Data was imputed using MAGIC (van Dijk et al., 2018) as recommended by the palantir documentation, with a step size of 1 (meaning data was imputed only using the very nearby cells). A starting cell for the trajectory (ID: TCATGTTGTTCTGACA) was identified using a high expression of CycE and D, and two terminal branch cells (IDs: AATCGACGTAATCAGA and AAGCGTTTCCTATTGT) were identified by choosing cells at the maximal points of the two major neural branches in the UMAP projections of the type-II cells. The choice of terminal cells was not necessary for the automatic identification of these 2 branches by palantir. They are provided here for data reproducibility purposes. Default parameters were used throughout the rest of the pseudotime trajectory inference.

## REFERENCES

Álvarez, J.-A., and Díaz-Benjumea, F.J. (2018). Origin and specification of type II neuroblasts in the Drosophila embryo. Development 145.

Bayraktar, O.A., and Doe, C.Q. (2013). Combinatorial temporal patterning in progenitors expands neural diversity. Nature 498, 449–455.

Bayraktar, O.A., Boone, J.Q., Drummond, M.L., and Doe, C.Q. (2010). Drosophila type II neuroblast lineages keep Prospero levels low to generate large clones that contribute to the adult brain central complex. Neural Dev. 5, 26.

Bello, B.C., Izergina, N., Caussinus, E., and Reichert, H. (2008). Amplification of neural stem cell proliferation by intermediate progenitor cells in Drosophila brain development. Neural Dev. 3, 5.

Boone, J.Q., and Doe, C.Q. (2008). Identification of Drosophila type II neuroblast lineages containing transit amplifying ganglion mother cells. Dev. Neurobiol. 68, 1185–1195.

Butler, A., Hoffman, P., Smibert, P., Papalexi, E., and Satija, R. (2018). Integrating single-cell transcriptomic data across different conditions, technologies, and species. Nat. Biotechnol. 36, 411–420.

Cao, J., Spielmann, M., Qiu, X., Huang, X., Ibrahim, D.M., Hill, A.J., Zhang, F., Mundlos, S., Christiansen, L., Steemers, F.J., et al. (2019). The single-cell transcriptional landscape of mammalian organogenesis. Nature 566, 496–502.

Choi, H.M.T., Schwarzkopf, M., Fornace, M.E., Acharya, A., Artavanis, G., Stegmaier, J., Cunha, A., and Pierce, N.A. (2018). Third-generation in situ hybridization chain reaction: multiplexed, quantitative, sensitive, versatile, robust. Development 145.

Cocanougher, B.T., Wittenbach, J.D., Long, X., Kohn, A.B., Norekian, T.P., Yan, J., Colonell, J., Masson, J.-B., Truman, J.W., Cardona, A., et al. (2019). Comparative single-cell transcriptomics of complete insect nervous systems. BioRxiv.

Deitcher, D.L., Ueda, A., Stewart, B.A., Burgess, R.W., Kidokoro, Y., and Schwarz, T.L. (1998). Distinct requirements for evoked and spontaneous release of neurotransmitter are revealed by mutations in the Drosophila gene neuronal-synaptobrevin. J. Neurosci. 18, 2028–2039.

Flybase curators (2019). FlyBase Reference Report: FlyBase, 2019-, Computation of D. melanogaster genes relevant to disease based on their orthology to human “disease genes”.

Gold, K.S., and Brand, A.H. (2014). Optix defines a neuroepithelial compartment in the optic lobe of the Drosophila brain. Neural Dev. 9, 18.

Greig, L.C., Woodworth, M.B., Galazo, M.J., Padmanabhan, H., and Macklis, J.D. (2013). Molecular logic of neocortical projection neuron specification, development and diversity. Nat. Rev. Neurosci. 14, 755–769.

Habib, N., Basu, A., Avraham-Davidi, I., Burks, T., Choudhury, S.R., Aguet, F., Gelfand, E., Ardlie, K., Weitz, D.A., Rozenblatt-Rosen, O., et al. (2017). DroNc-Seq: Deciphering cell types in human archived brain tissues by massively-parallel single nucleus RNA-seq. BioRxiv.

Han, X., Wang, R., Zhou, Y., Fei, L., Sun, H., Lai, S., Saadatpour, A., Zhou, Z., Chen, H., Ye, F., et al. (2018). Mapping the Mouse Cell Atlas by Microwell-Seq. Cell 172, 1091-1107.e17.

Homem, C.C.F., and Knoblich, J.A. (2012). Drosophila neuroblasts: a model for stem cell biology. Development 139, 4297–4310.

Huang, D.W., Sherman, B.T., and Lempicki, R.A. (2009a). Bioinformatics enrichment tools: paths toward the comprehensive functional analysis of large gene lists. Nucleic Acids Res. 37, 1–13.

Huang, D.W., Sherman, B.T., and Lempicki, R.A. (2009b). Systematic and integrative analysis of large gene lists using DAVID bioinformatics resources. Nat. Protoc. 4, 44–57.

Kalhor, R., Kalhor, K., Mejia, L., Leeper, K., Graveline, A., Mali, P., and Church, G.M. (2018). Developmental barcoding of whole mouse via homing CRISPR. Science 361.

Kiselev, V.Y., Andrews, T.S., and Hemberg, M. (2019). Challenges in unsupervised clustering of single-cell RNA-seq data. Nat. Rev. Genet. 20, 273–282.

Kose, H., Rose, D., Zhu, X., and Chiba, A. (1997). Homophilic synaptic target recognition mediated by immunoglobulin-like cell adhesion molecule Fasciclin III. Development 124, 4143–4152.

Kucherenko, M.M., Ilangovan, V., Herzig, B., Shcherbata, H.R., and Bringmann, H. (2016). TfAP-2 is required for night sleep in Drosophila. BMC Neurosci. 17, 72.

Lai, E.C., Bodner, R., and Posakony, J.W. (2000). The enhancer of split complex of Drosophila includes four Notch-regulated members of the bearded gene family. Development 127, 3441–3455.

Landskron, L., Steinmann, V., Bonnay, F., Burkard, T.R., Steinmann, J., Reichardt, I., Harzer, H., Laurenson, A.-S., Reichert, H., and Knoblich, J.A. (2018). The asymmetrically segregating lncRNA cherub is required for transforming stem cells into malignant cells. Elife 7.

Lane, M.E., Sauer, K., Wallace, K., Jan, Y.N., Lehner, C.F., and Vaessin, H. (1996). Dacapo, a cyclin-dependent kinase inhibitor, stops cell proliferation during Drosophila development. Cell 87, 1225–1235.

Lin, Y., Ghazanfar, S., Wang, K.Y.X., Gagnon-Bartsch, J.A., Lo, K.K., Su, X., Han, Z.-G., Ormerod, J.T., Speed, T.P., Yang, P., et al. (2019). scMerge leverages factor analysis, stable expression, and pseudoreplication to merge multiple single-cell RNA-seq datasets. Proc Natl Acad Sci USA 116, 9775–9784.

Liu, K.S.Y., Siebert, M., Mertel, S., Knoche, E., Wegener, S., Wichmann, C., Matkovic, T., Muhammad, K., Depner, H., Mettke, C., et al. (2011). RIM-binding protein, a central part of the active zone, is essential for neurotransmitter release. Science 334, 1565–1569.

Li, X., Chen, R., and Zhu, S. (2017). bHLH-O proteins balance the self-renewal and differentiation of Drosophila neural stem cells by regulating Earmuff expression. Dev. Biol. 431, 239–251.

Li, Y., Walker, L.A., Zhao, Y., Edwards, E.M., Michki, N.S., Cheng, H.P.J., Ghazzi, M., Chen, T.Y., Chen, M., Roossien, D.H., et al. (2020). Bitbow: a digital format of Brainbow enables highly efficient neuronal lineage tracing and morphology reconstruction in single brains. BioRxiv.

Macosko, E.Z., Basu, A., Satija, R., Nemesh, J., Shekhar, K., Goldman, M., Tirosh, I., Bialas, A.R., Kamitaki, N., Martersteck, E.M., et al. (2015). Highly Parallel Genome-wide Expression Profiling of Individual Cells Using Nanoliter Droplets. Cell 161, 1202–1214.

Maier, D., Marte, B.M., Schäfer, W., Yu, Y., and Preiss, A. (1993). Drosophila evolution challenges postulated redundancy in the E(spl) gene complex. Proc Natl Acad Sci USA 90, 5464–5468.

Mariano, V., Achsel, T., Bagni, C., and Kanellopoulos, A.K. (2020). Modelling learning and memory in Drosophila to understand Intellectual Disabilities. Neuroscience.

Monastirioti, M., Giagtzoglou, N., Koumbanakis, K.A., Zacharioudaki, E., Deligiannaki, M., Wech, I., Almeida, M., Preiss, A., Bray, S., and Delidakis, C. (2010). Drosophila Hey is a target of Notch in asymmetric divisions during embryonic and larval neurogenesis. Development 137, 191–201.

Monge, I., Krishnamurthy, R., Sims, D., Hirth, F., Spengler, M., Kammermeier, L., Reichert, H., and Mitchell, P.J. (2001). Drosophila transcription factor AP-2 in proboscis, leg and brain central complex development. Development 128, 1239–1252.

Mukai, M., Hayashi, Y., Kitadate, Y., Shigenobu, S., Arita, K., and Kobayashi, S. (2007). MAMO, a maternal BTB/POZ-Zn-finger protein enriched in germline progenitors is required for the production of functional eggs in Drosophila. Mech. Dev. 124, 570–583.

Müller, M., Genç, Ö., and Davis, G.W. (2015). RIM-binding protein links synaptic homeostasis to the stabilization and replenishment of high release probability vesicles. Neuron 85, 1056–1069.

de Nooij, J.C., Letendre, M.A., and Hariharan, I.K. (1996). A cyclin-dependent kinase inhibitor, Dacapo, is necessary for timely exit from the cell cycle during Drosophila embryogenesis. Cell 87, 1237–1247.

Ntranos, V., Yi, L., Melsted, P., and Pachter, L. (2018). Identification of transcriptional signatures for cell types from single-cell RNA-Seq. BioRxiv.

Ogawa, L.M., and Vallender, E.J. (2014). Evolutionary conservation in genes underlying human psychiatric disorders. Front. Hum. Neurosci. 8, 283.

Pei, W., Feyerabend, T.B., Rössler, J., Wang, X., Postrach, D., Busch, K., Rode, I., Klapproth, K., Dietlein, N., Quedenau, C., et al. (2017). Polylox barcoding reveals haematopoietic stem cell fates realized in vivo. Nature 548, 456–460.

Ponti, G., Obernier, K., Guinto, C., Jose, L., Bonfanti, L., and Alvarez-Buylla, A. (2013). Cell cycle and lineage progression of neural progenitors in the ventricular-subventricular zones of adult mice. Proc Natl Acad Sci USA 110, E1045–54.

Qiu, X., Mao, Q., Tang, Y., Wang, L., Chawla, R., Pliner, H.A., and Trapnell, C. (2017). Reversed graph embedding resolves complex single-cell trajectories. Nat. Methods 14, 979–982.

Raj, B., Wagner, D.E., McKenna, A., Pandey, S., Klein, A.M., Shendure, J., Gagnon, J.A., and Schier, A.F. (2017). Simultaneous single-cell profiling of lineages and cell types in the vertebrate brain by scGESTALT. BioRxiv.

Ren, J., Isakova, A., Friedmann, D., Zeng, J., Grutzner, S.M., Pun, A., Zhao, G.Q., Kolluru, S.S., Wang, R., Lin, R., et al. (2019). Single-cell transcriptomes and whole-brain projections of serotonin neurons in the mouse dorsal and median raphe nuclei. Elife 8.

Ren, Q., Yang, C.-P., Liu, Z., Sugino, K., Mok, K., He, Y., Ito, M., Nern, A., Otsuna, H., and Lee, T. (2017). Stem Cell-Intrinsic, Seven-up-Triggered Temporal Factor Gradients Diversify Intermediate Neural Progenitors. Curr. Biol. 27, 1303–1313.

Saunders, A., Macosko, E., Wysoker, A., Goldman, M., Krienen, F., Bien, E., Baum, M., Wang, S., Goeva, A., Nemesh, J., et al. (2018). A Single-Cell Atlas of Cell Types, States, and Other Transcriptional Patterns from Nine Regions of the Adult Mouse Brain. BioRxiv.

Schinaman, J.M., Giesey, R.L., Mizutani, C.M., Lukacsovich, T., and Sousa-Neves, R. (2014). The KRÜPPEL-like transcription factor DATILÓGRAFO is required in specific cholinergic neurons for sexual receptivity in Drosophila females. PLoS Biol. 12, e1001964.

Setty, M., Kiseliovas, V., Levine, J., Gayoso, A., Mazutis, L., and Pe’er, D. (2019). Characterization of cell fate probabilities in single-cell data with Palantir. Nat. Biotechnol. 37, 451–460.

Snow, P.M., Bieber, A.J., and Goodman, C.S. (1989). Fasciclin III: a novel homophilic adhesion molecule in Drosophila. Cell 59, 313–323.

Soldatov, R., Kaucka, M., Kastriti, M.E., Petersen, J., Chontorotzea, T., Englmaier, L., Akkuratova, N., Yang, Y., Häring, M., Dyachuk, V., et al. (2019). Spatiotemporal structure of cell fate decisions in murine neural crest. Science 364.

Spanjaard, B., Hu, B., Mitic, N., Olivares-Chauvet, P., Janjuha, S., Ninov, N., and Junker, J.P. (2018). Simultaneous lineage tracing and cell-type identification using CRISPR-Cas9-induced genetic scars. Nat. Biotechnol. 36, 469–473.

Syed, M.H., Mark, B., and Doe, C.Q. (2017). Steroid hormone induction of temporal gene expression in Drosophila brain neuroblasts generates neuronal and glial diversity. Elife 6.

Traag, V.A., Waltman, L., and van Eck, N.J. (2019). From Louvain to Leiden: guaranteeing well-connected communities. Sci. Rep. 9, 5233.

Tran, T.N., and Bader, G. (2019). Tempora: cell trajectory inference using time-series single-cell RNA sequencing data. BioRxiv.

Tumbar, T., Guasch, G., Greco, V., Blanpain, C., Lowry, W.E., Rendl, M., and Fuchs, E. (2004). Defining the epithelial stem cell niche in skin. Science 303, 359–363.

Turek, M., Lewandrowski, I., and Bringmann, H. (2013). An AP2 transcription factor is required for a sleep-active neuron to induce sleep-like quiescence in C. elegans. Curr. Biol. 23, 2215–2223.

Veling, M.W., Li, Y., Veling, M.T., Litts, C., Michki, N., Liu, H., Ye, B., and Cai, D. (2019). Identification of Neuronal Lineages in the Drosophila Peripheral Nervous System with a “Digital” Multi-spectral Lineage Tracing System. Cell Rep. 29, 3303-3312.e3.

Weber, T.S., Dukes, M., Miles, D.C., Glaser, S.P., Naik, S.H., and Duffy, K.R. (2016). Site-specific recombinatorics: in situ cellular barcoding with the Cre Lox system. BMC Syst. Biol. 10, 43.

Wells, R.E., Barry, J.D., Warrington, S.J., Cuhlmann, S., Evans, P., Huber, W., Strutt, D., and Zeidler, M.P. (2013). Control of tissue morphology by Fasciclin III-mediated intercellular adhesion. Development 140, 3858–3868.

Weng, M., Golden, K.L., and Lee, C.-Y. (2010). dFezf/Earmuff maintains the restricted developmental potential of intermediate neural progenitors in Drosophila. Dev. Cell 18, 126–135.

Wolf, F.A., Angerer, P., and Theis, F.J. (2018). SCANPY: large-scale single-cell gene expression data analysis. Genome Biol. 19, 15.

Yang, L., Titlow, J., Ennis, D., Smith, C., Mitchell, J., Young, F.L., Waddell, S., Ish-Horowicz, D., and Davis, I. (2017). Single molecule fluorescence in situ hybridisation for quantitating post-transcriptional regulation in Drosophila brains. Methods 126, 166–176.

Zhong, S., Zhang, S., Fan, X., Wu, Q., Yan, L., Dong, J., Zhang, H., Li, L., Sun, L., Pan, N., et al. (2018). A single-cell RNA-seq survey of the developmental landscape of the human prefrontal cortex. Nature 555, 524–528.

Zhou, Q., Liu, M., Xia, X., Gong, T., Feng, J., Liu, W., Liu, Y., Zhen, B., Wang, Y., Ding, C., et al. (2017). A mouse tissue transcription factor atlas. Nat. Commun. 8, 15089.

Ziegenhain, C., Vieth, B., Parekh, S., Reinius, B., Guillaumet-Adkins, A., Smets, M., Leonhardt, H., Heyn, H., Hellmann, I., and Enard, W. (2017). Comparative Analysis of Single-Cell RNA Sequencing Methods. Mol. Cell 65, 631-643.e4.

