## Supplementary material for "The molecular landscape of neural differentiation in the developing *Drosophila* brain revealed by targeted scRNA-seq and a multi-informatic analysis paradigm": Resource Table

### Key Resources table

| REAGENT or RESOURCE | SOURCE | IDENTIFIER |
|---|---|---|
| **Antibodies** | | |
| Mouse monoclonal anti-Fas3 | DHSB | Cat# 7G10, RRID:AB_528238 |
| Rat monoclonal anti-Dpn | Lee, C.Y., Robinson, K.J., and Doe, C.Q. (Nature, 2006a) | NA |
| Donkey-anti-Ms (AF488) | Jackson ImmunoResearch Laboratories, Inc. | Cat# 715-545-151 |
| Donkey-anti-Rt (AF647) | Jackson ImmunoResearch Laboratories, Inc. | Cat# 712-605-150 |
| **Chemicals, Peptides, and Recombinant Proteins** | | |
| Papain | Millipore Sigma | Cat# P4762-25MG |
| Collagenase type I | Millipore Sigma | Cat# SCR103 |
| E-64 | Millipore Sigma | Cat# E3132-1MG |
| Fetal Bovine Serum | Millipore Sigma | Cat# F0926-50ML |
| Schneider's Media | Millipore Sigma | Cat# S0146-500ML |
| DRAQ5 | abcam | Cat# ab108410 |
| Dextran sulfate, 50% solution | Millipore Sigma | Cat# S4031 |
| **Critical Commercial Assays** | | |
| 10X chromium v3 single-cell gene expression kit | 10X Genomics | Cat# 1000154 |
| **Deposited Data** | | |
| Raw and analyzed data | This study | GEO: [ID here] |

| Experimental Models: Organisms/Strains | | |
|---|---|---|
| D. melanogaster, R9D11-Gal4 driver line: w[1118]; P{y[+t7.7] w[+mC]=GMR9D11-GAL4}attP2 | BDSC | RRID:BDSC_40731 |
| D. melanogaster, R9D11-CD4::tdTomato membrane reporter line: w[1118]; P{y[+t7.7] w[+mC]=R9D11-CD4-tdTom}attP2/TM6B, Tb[1] | BDSC | RRID:BDSC_40731 |
| D. melanogaster: yw;;UAS-hH2B::2xmNG | This study | NA |
| D. melanogaster: yw;;UAS-hH2B::2xTagBFP2 | This study | NA |
| D. melanogaster, Sp1::EGFP protein fusion reporter line: w[1118]; PBac{y[+mDint2] w[+mC]=Sp1-EGFP.S}VK00033 | BDSC | RRID:BDSC_38669 |
| D. melanogaster, UAS-IVS-myr::tdTomato membrane reporter line: w[*]; P{y[+t7.7] w[+mC]=10XUAS-IVS-myr::tdTomato}attP40 | BDSC | RRID:BDSC_32222 |
| **Oligonucleotides** | | |
| *mNeonGreen* HCR probe set | Molecular Instruments | PRC014 |
| *CycE* HCR probe set | Molecular Instruments | PRD167 |
| *D* HCR probe set | Molecular Instruments | PRC881 |
| *Sp1* HCR probe set | Molecular Instruments | PRC883 |
| *TfAP-2* HCR probe set | Molecular Instruments | PRD168 |
| *Fas3* HCR probe set | Molecular Instruments | PRC900 |
| *ytr* HCR probe set | Molecular Instruments | PRE680 |
| *E(spl)m6-BFM* HCR probe set | Molecular Instruments | PRE684 |
| *tap* HCR probe set | Molecular Instruments | PRE682 |
| *jim* HCR probe set | Molecular Instruments | PRE686 |
| **Software and Algorithms** | | |
| Fiji/ImageJ | Schindelin et al., 2012 | https://fiji.sc/ |
| scanpy scRNA-seq analysis software | Wolf, F., Angerer, P. & Theis, F., 2018 | RRID:SCR_018139 |
| Fiji/ImageJ | Schindelin et al., 2012 | https://fiji.sc/ |
| STAR RNA-seq aligner | Dobin et al., 2013 | RRID:SCR_015899 https://github.com/alexdobin/STAR |
| MiCV web tool | This study | https://micv.works https://github.com/cailabumich/MiCV |
