## Supplementary Figures for "The molecular landscape of neural differentiation in the developing *Drosophila* brain revealed by targeted scRNA-seq and a multi-informatic analysis paradigm"

### SUPPLEMENTAL INFORMATION

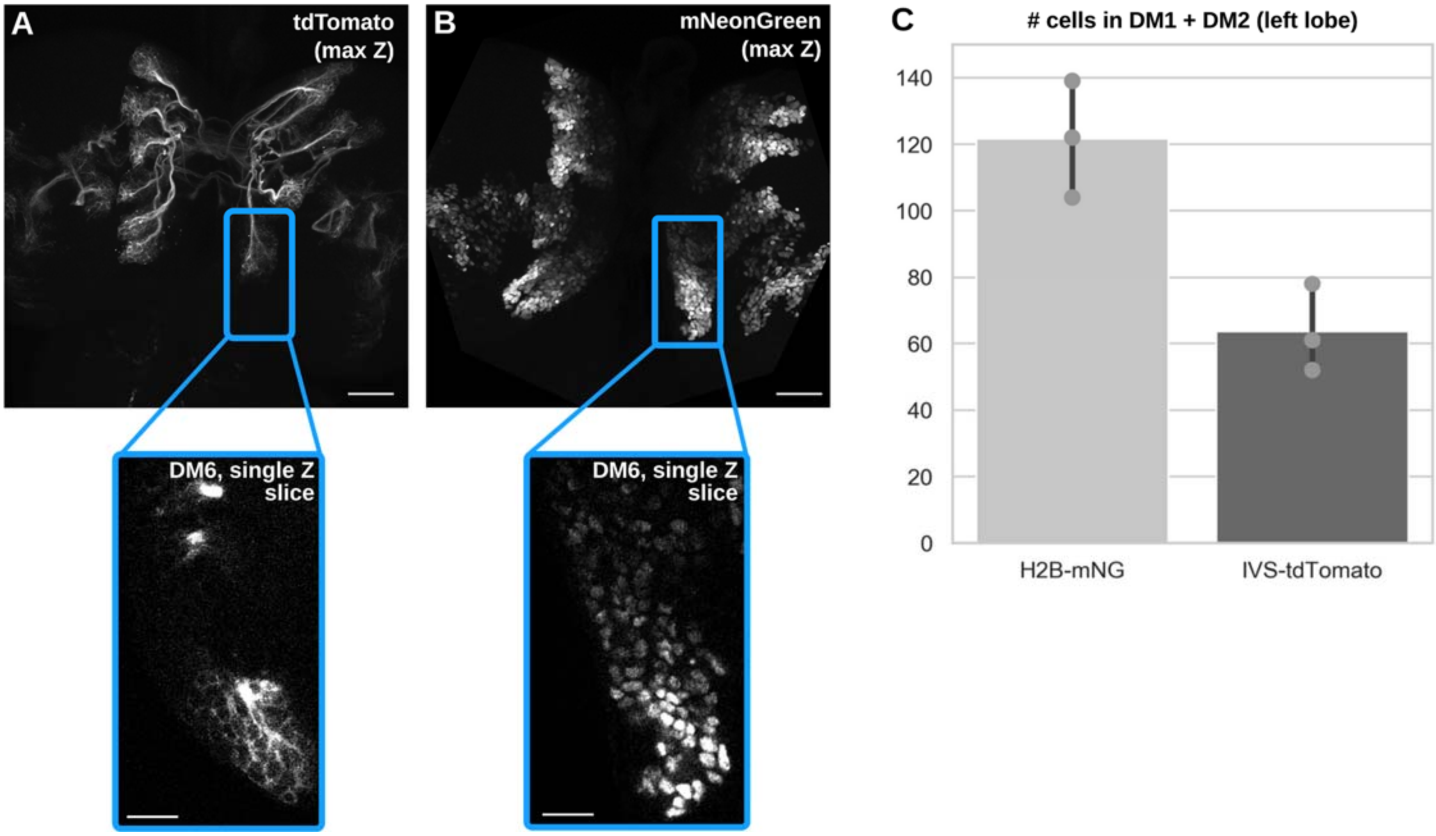


**Supplemental Figure 1. A novel, long-lasting nucleus UAS-hH2B::2xmNeonGreen (mNG) reporter labels more cells in the type-II progenies than the membrane UAS-IVS-myc::tdTomato reporter.**
**(A)** The membrane-bound tdTomato reporter is driven under the control of R9D11-Gal4 and its lineage labeling is compared to that of our **(B)** nucleus-targeted 2xmNG reporter. **(C)** Quantifications of labeled cells in clusters DM1 and DM2 in late third instar larvae brains. Scale bars, 30 μm in overviews of (A,B), 10 μm in insets of (A,B).

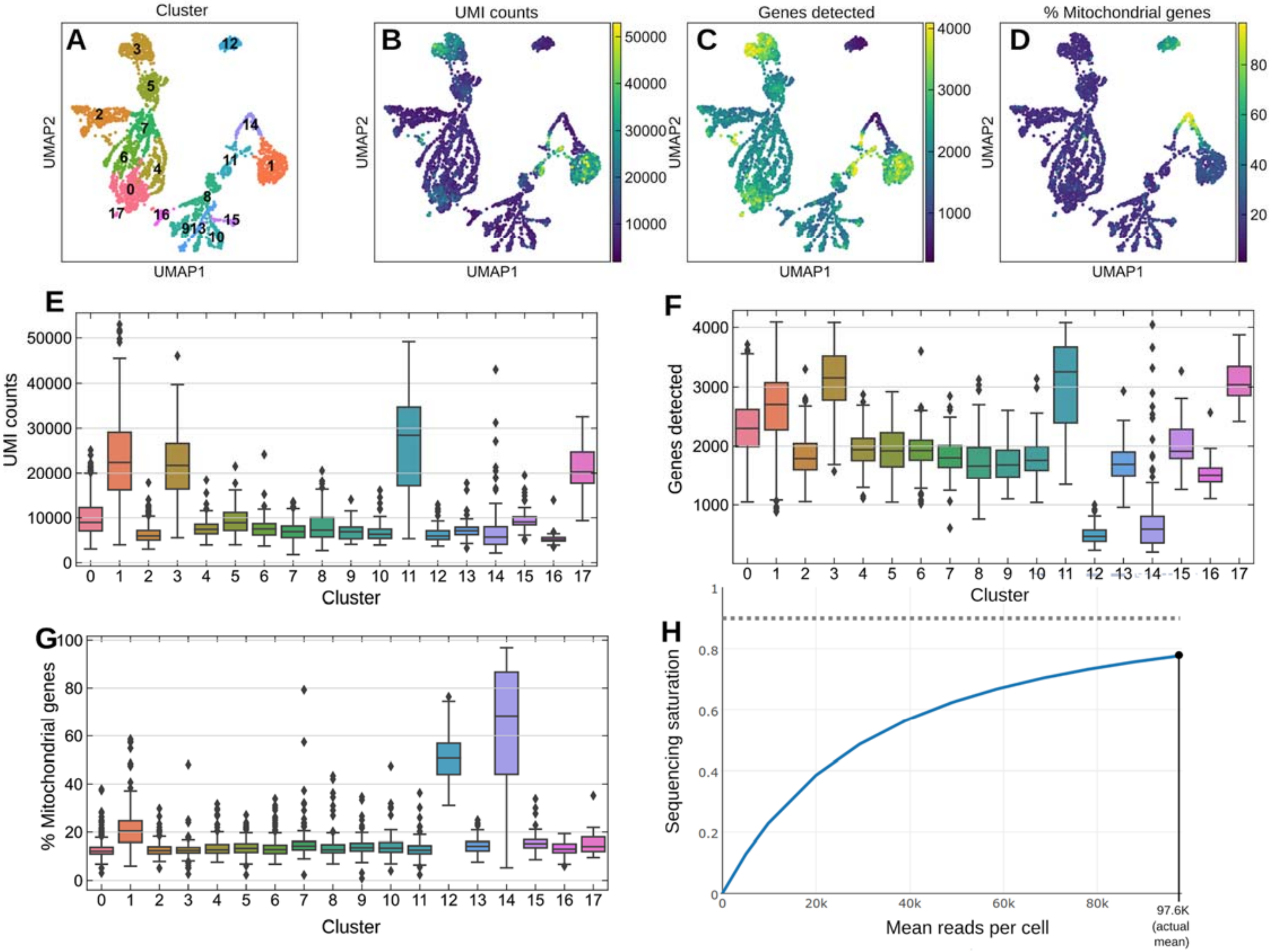


**Supplemental Figure 2. Sequencing QC metrics indicate that captured cells are healthy and diverse in transcriptional activity.** **(A)** UMAP with colorimetric and numerical labels for each automatically assigned cluster (Leiden algorithm, resolution=0.5). **(B-D)** UMAP of cells overlaid with their number of UMI counts, number of unique genes detected, and percentage of mitochondrial genes, respectively. **(E-G)** the QC metrics from above but summarized as boxplots on a per-cluster basis. Clusters 1, 3, and 11 are large groups of cells that have particularly high UMI counts and gene detection rates, indicating that they are transcriptionally very active. Cluster 3 is the group of type-II derived INPs described in this work. Cluster 1 cells are likely glia based on the expression of *repo* (data not shown) and cluster 11 cells are likely progenitors in the OL cells based on the expression of *CycE* (data not shown). Cluster 0 is a group of maturing neurons that has a higher than average gene detection rate, and strongly expresses *Imp* (data not shown), an IGF-II RNA-binding protein that is responsible for a number of RNA trafficking functions, notably being required for axonal growth and remodeling (Medioni et al. 2014). As the type-II neuronal progenies extend large axonal bundles across the commissure during development, it is possible that this transcriptionally active group of *Imp*+ neurons are the ones actively undergoing this process. **(H)** Predicted sequencing saturation curve generated using CellRanger, indicating that at our sequencing depth we have recovered nearly 80% of unique genes that might be found in each cell.

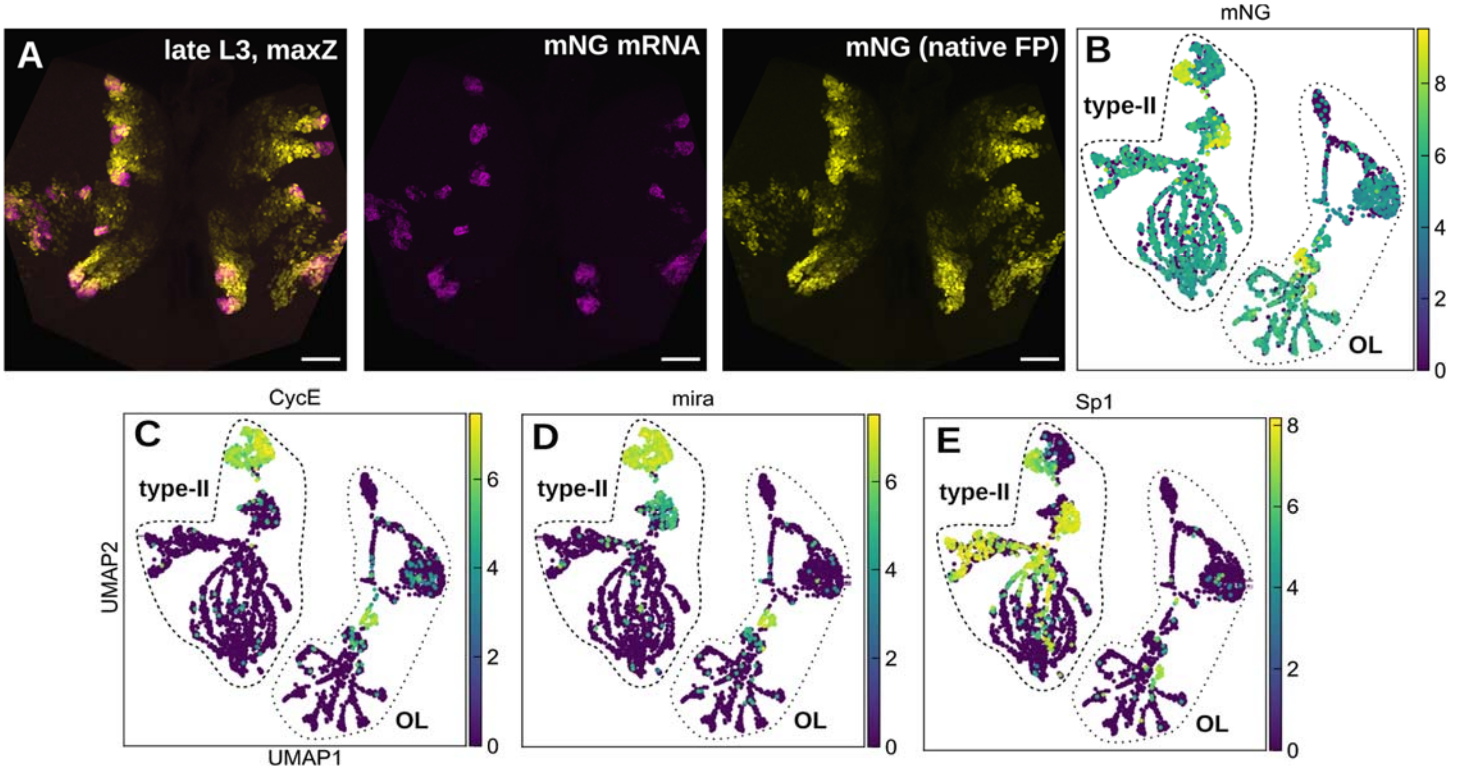


**Supplemental Figure 3. R9D11-Gal4 driven reporter mRNA expression is restricted to a small portion of each type-II lineage.** **(A)** A composite maximum z-projection of a late third instar larval brain expressing our novel UAS-hH2B::2xmNeonGreen (mNG) reporter under the control of R9D11-Gal4, with native mNG fluorescence labeling the type-II progenies and mNG mRNA labeled using our mNG HCR v3 probes. At the tip of each type-II lineage, there is a burst of expression of mNG transcripts (middle panel) that does *not* persist throughout the lineage but rather remains restricted to what is presumably the youngest mINPs. **(B)** This assessment is further validated using our scRNA-seq data, wherein we find that mapped mNG transcript expression is multiple log2-fold higher in young mINPs and their daughter GMCs, based on the expression of *CycE/mira* for mINPs and *Sp1* for *young* mINPs and their progeny **(C-E)** (see also **Fig. 3** in the main text). In the optic lobe (the large connected group of cells on the right-hand side of the UMAP projection), the expression is not restricted to cells with the highest *CycE/mira* expression and so it is possible that the R9D11 enhancer element is active in a non-progenitor population in the optic lobe. Scale bars: 30 μm.

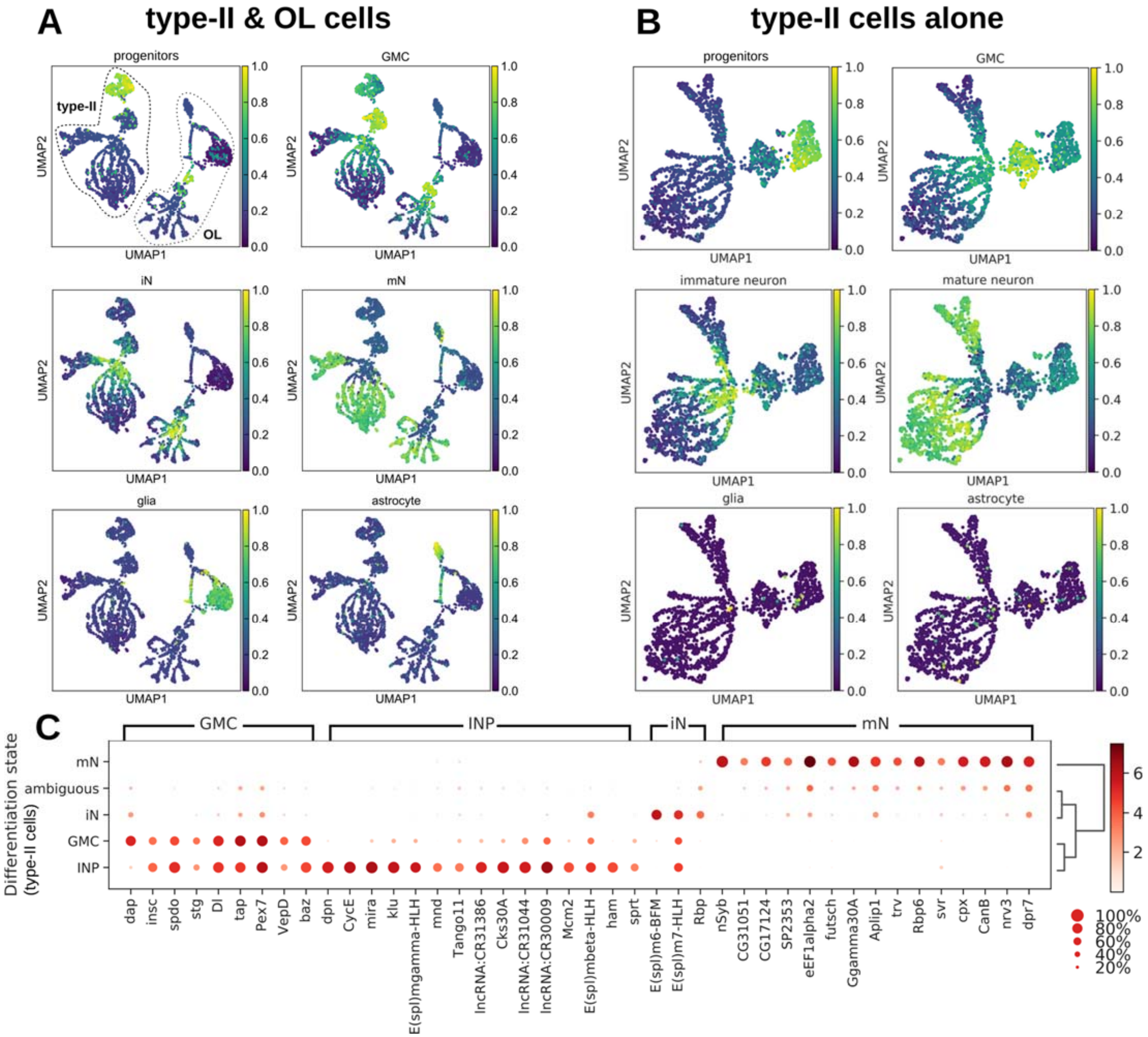


**Supplemental Figure 4. Marker gene-based differentiation state scoring enables robust identification of cell differentiation state without manual annotation.**

**(A)** All cells were scored using the score_genes function from scanpy with the following marker genes defining each differentiation state (see table below). These scores were normalized to be within the range of [0,1], with 1 indicating that all of the marker genes for that cell type were expressed at high levels in that particular cell. These genes were largely selected based on literature references (listed in the table below), but some were identified in this and other very recent scRNA-seq works. **(B)** Type-II NB derived cells were scored as described in (A). **(C)** Marker gene analysis revealed genes that specify clusters of cells in distinct maturation/differentiation states. To be noted that many GMC marker genes are also highly expressed in INPs. Although pseudotime analysis provides a more holistic view of a gene's dynamic change along the cell differentiation trajectory (**Fig. 2**), this small set of genes are robust identifiers for determining cell differentiation states. INP, intermediate progenitor cell; GMC, ganglion mother cell; iN, immature neuron; mN, matured/maturing neuron.

| progenitors | GMCs | immature neurons | mature neurons | glia | astrocytes |
|---|---|---|---|---|---|
| *CycE, mira, dpn* | *insb, insc, spdo, nerfin-1, dap* | *Hey, E(spl)m6-BFM* | *nSyb, lncRNA:noe, jim* | *repo, gcm* | *Gat, alrm* |

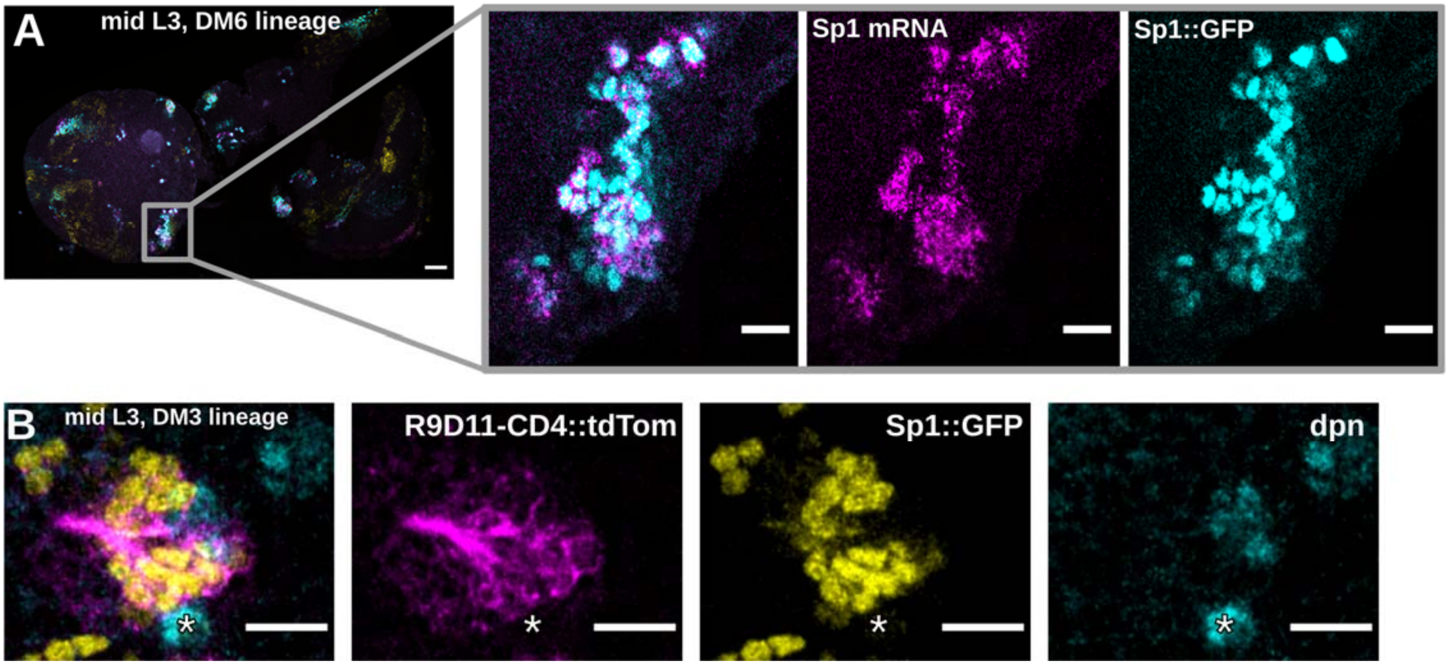


**Supplemental Figure 5. Sp1::GFP fusion protein and Sp1 mRNA co-localize in situ and label both dpn+ INPs and axon-producing neurons.**

**(A)** Single z-slice of a mid 3rd instar Sp1::GFP transgenic larval brain, showing native fluorescence of GFP in cyan (right), HCR stained *Sp1* mRNA in magenta (middle), and a composite of the two (left). *Sp1* mRNA signal is made up of puncta scattered around the labeled cells. *Sp1* protein, being a transcription factor, leads to GFP expression being largely localized to the nucleus. **(B)** Single z-slice of a mid 3rd instar ;R9D11-CD4::tdTomato;Sp1::GFP larval brain stained with an antibody specific to dpn. The co-localization of dpn, Sp1, and membrane-bound tdTomato (mT) proteins indicates that Sp1 is translated in both neurons and INPs of the type-II lineage, as evidenced by the labelling of cells that either produce mT-labeled axons or are dpn+, respectively. Scale bars: 30 µm in the overview of (A); 10 µm in insets of (B) and (B).

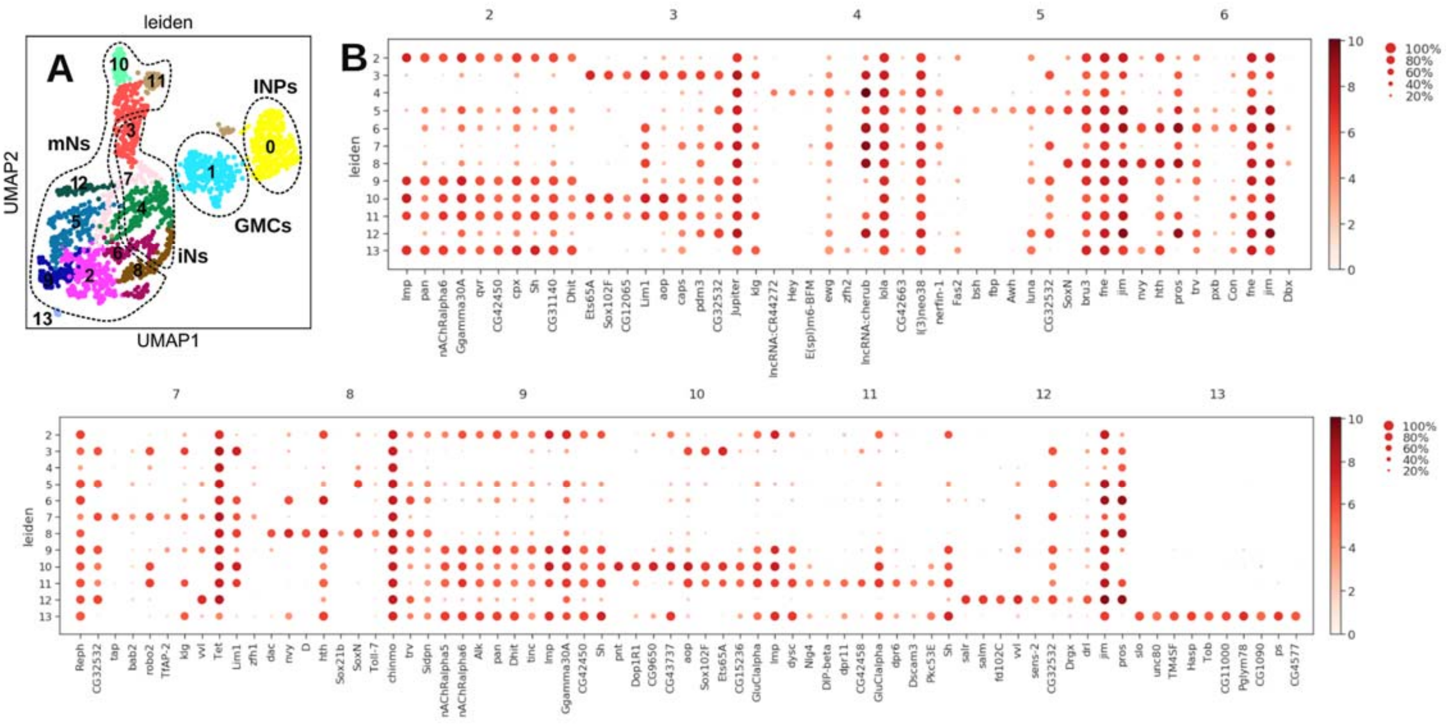


**Supplemental Figure 6. Marker genes of subtype-specific immature/maturing neurons.**
**(A)** UMAP plot with automatic cluster assignments (resolution = 0.6). Dotted outlines indicate groups of neurons, whose terminal fates are specified by the transcription factors *Sp1*, *bsh*, and *D* as described in (**Fig. 5**). **(B)** The top 10 marker genes identified for each of the neural clusters in this scRNA-seq dataset.
